# Unique functions of two overlapping *PAX6* retinal enhancers

**DOI:** 10.1101/2022.11.25.517987

**Authors:** Kirsty Uttley, Andrew S. Papanastasiou, Manuela Lahne, Jennifer M. Brisbane, Ryan B. MacDonald, Wendy A. Bickmore, Shipra Bhatia

## Abstract

Enhancers play a critical role in development by precisely modulating spatial, temporal, and cell type-specific gene expression. Sequence variants in enhancers have been implicated in disease, however establishing the functional consequences of these variants is challenging due to a lack of understanding of precise cell types and developmental stages where the enhancers are normally active. *PAX6* is the master regulator of eye development, and has a regulatory landscape containing multiple enhancers driving expression in the eye. Whether these enhancers perform additive, redundant, or distinct functions is unknown. Here we describe the precise cell types and regulatory activity of two *PAX6* retinal enhancers, HS5 and NRE. Using a unique combination of live imaging and single-cell RNA sequencing in dual enhancer-reporter zebrafish embryos, we find significant differences in the spatiotemporal activity of these enhancers, and show that HS5 and NRE are active in distinct cell types of the developing retina. Our results show that although overlapping, these enhancers have distinct activities in different cell types and therefore likely non-redundant functions. This work demonstrates that unique cell type-specific functions can be uncovered for apparently similar enhancers when investigated at high resolution in vivo.

## Introduction

Regulatory elements such as enhancers control the activation of target genes in precise spatial and temporal patterns, ensuring proper gene expression and the successful development of complex organisms (Long et al., 2016). The human genome contains millions of predicted enhancers, and non-coding mutations affecting enhancers can cause Mendelian disease, as well as contribute to complex phenotypes and drive evolutionary differences between species (Lettice et al., 2003; Consortium, 2012; Maurano et al., 2012; Smemo et al., 2012; Bhatia et al., 2013; Long et al., 2020). Unravelling the functions of non-coding elements is key to understanding the regulatory rules of enhancers, and the consequences of sequence variation. However, understanding the functions of enhancers remains a fundamental challenge (Meuleman et al., 2020; Jindal and Farley, 2021). In contrast to protein-coding sequences, it is not possible to predict the function of an enhancer from sequence alone. Transcription factor binding sites within enhancers often correspond to suboptimal binding motifs, making it hard to predict which binding sites and transcription factors are important for function (Farley et al., 2015, 2016). Another confounding problem is the cooperativity, or perhaps redundancy, of enhancers. Several studies have shown that mutation or loss of an enhancer does not necessarily cause observable, gross phenotypes in animal models (Osterwalder et al., 2018; Kvon et al., 2020; Snetkova et al., 2021). This may be due to the buffering of enhancer loss by multiple elements driving similar patterns of gene expression, acting redundantly. Such overlapping enhancers can however have unique or additive effects, but the phenotypes arising from mutation of these elements can be subtle and highly cell type specific (Dickel et al., 2018; Long et al., 2020). It is therefore important to understand the precise functions of enhancers, particularly for elements with similar tissue-specific domains of activity.

The expression of pleiotropic developmental genes is controlled by multiple tissue-specific enhancers, and it is common for such genes to have multiple elements with apparently similar spatiotemporal activities (Kvon et al., 2021). An example is the regulatory landscape of *PAX6*, encoding a developmental transcription factor which, among other functions, is a master regulator of eye development, controlling functions ranging from the specification of the eye field, to maintaining progenitor populations, and influencing the differentiation of multiple cell types (van Heyningen, 2002). The large *PAX6* regulatory landscape contains several identified enhancers active in overlapping domains of the developing lens and retina (Kammandel et al., 1999; Kleinjan et al., 2001; McBride et al., 2011; Ravi et al., 2013; Bhatia et al., 2014; Lima Cunha et al., 2019) (Figure 1A). The sequence of *PAX6* is highly conserved from flies to humans, as is its role in the specification of eye development (Halder et al., 1995; Onuma et al., 2002). Human *PAX6* regulatory elements, and their activities, also show high levels of conservation in vertebrate genomes (Williams et al., 1998; Griffin et al., 2002; Tyas et al., 2006; Bhatia et al., 2014). Mutations affecting these *PAX6* regulatory elements, ranging from large scale genome rearrangements to single nucleotide point mutations, can cause eye malformations such as aniridia (Lauderdale et al., 2000; Kleinjan et al., 2001; Bhatia et al., 2013). As is the case for other developmental loci, understanding the functions of individual *PAX6* enhancer elements, and whether they act in redundant, additive, or distinct ways, will help to decode non-coding mutations at this locus and further our understanding of the mechanisms of enhancer activity during development.

**Figure 1.**
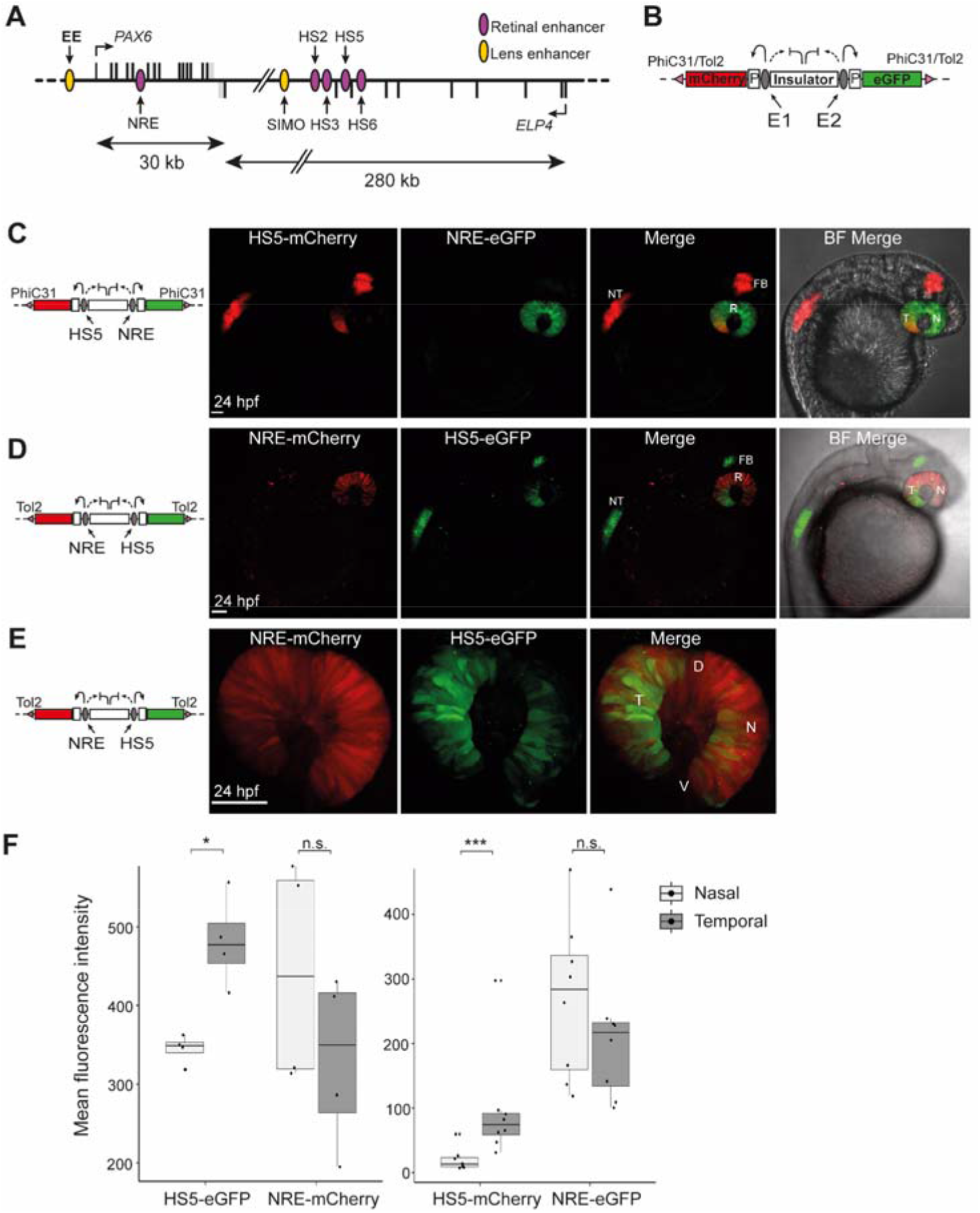
Activity of HS5 and NRE in a dual enhancer-reporter system during zebrafish embryonic development. **(A)** Map of human *PAX6* regulatory locus showing the position of eye enhancers including HS5 and NRE (retinal, purple; lens, yellow). **(B)** Dual enhancer-reporter injection construct. mCherry and eGFP are transcribed from a minimal *gata2* promoter (P) activated by enhancer E1 or E2. An insulator based on the chicken HS4 sequence separates the enhancers. Targeted PhiC31 integration or random Tol2 integration is used to insert the dual-reporter construct into the zebrafish genome (Bhatia et al., 2021). **(C)** Live imaging of a 24 hpf NRE-eGFP/HS5-mCherry F1 embryo (10x objective). NRE (eGFP) is active throughout the retina (R). HS5 (mCherry) is active in the forebrain (FB), neural tube (NT) and in the retina where activity is highest in the temporal (T) half of the retina, compared to the nasal (N) side. **(D)** Live imaging of a 24 hpf NRE-mCherry/HS5-eGFP F1 embryo showing the activity of HS5 (eGFP) in FB, NT, and predominantly the temporal retina, towards the ventral (V) side as opposed to dorsal (D). NRE (mCherry) is active throughout the retina (10x objective). **(E)** As in **(D)** but at higher resolution (40x water immersion objective). Scale bars 50 µm. **(F)** Quantification of mean fluorescence intensity for mCherry and eGFP in the nasal versus temporal retina at 24 hpf in NRE-mCherry/HS5-eGFP (left) and NRE-eGFP/HS5-mCherry (right) F1 embryos. The activity of HS5 (eGFP or mCherry), is significantly higher in the temporal retina. n F1 embryos imaged ≥ 4. Wilcoxon test results: ns, not significant; *, p<0.05; ***, p<0.001. Scale bars 50 µm.

Here we use a previously developed dual enhancer-reporter assay in *Danio rerio* (zebrafish) to dissect the activity of two overlapping human *PAX6* retinal enhancers, HS5 and NRE (Bhatia et al., 2021). This assay has previously been used to study the activity of human enhancers, and recapitulates known patterns of activity for wild type and mutant sequences (Bhatia et al., 2021). Compared to enhancer-reporter assays in cell lines, this assay preserves the tissue-specific context of enhancer activities and allows visualisation of their spatial and temporal domains of activity in live animals during development, and in-depth characterisation and comparison of two enhancers in the same embryo. Using a unique combination of live imaging and single-cell RNA sequencing (scRNA-seq) we uncover differences in the spatial, temporal, and cell type activities of HS5 and NRE. Our results demonstrate how distinct differences between two enhancers with overlapping activities can be revealed by high-resolution in vivo analysis.

## Results

### HS5 and NRE have unique patterns of spatial and temporal activity in the developing retina

HS5 and NRE are two human *PAX6* enhancers (Figure 1A) known to be active in the developing retina, with apparent overlap in their domains of activity (Plaza et al., 1995; Kammandel et al., 1999; McBride et al., 2011). The activity of these elements has been assessed individually in enhancer-reporter assays, using highly conserved sequences from mouse, quail, lamprey and elephant shark (Plaza et al., 1995; Kammandel et al., 1999; McBride et al., 2011; Ravi et al., 2013, 2019). Studies on the mouse NRE sequence (also referred to as α-enhancer) have indicated that NRE is active in retinal progenitors and amacrine cell development (Marquardt et al., 2001; Kim et al., 2017; Dupacova et al., 2021). However, the precise stage and cell type-specific functions of these enhancers have not been fully characterised. In order to define and directly compare the functional activities of HS5 and NRE we created a dual enhancer-reporter zebrafish line using the QSTARZ system (Figure 1B) (Bhatia et al., 2021). This line contains a dual enhancer-reporter cassette, with the enhancers separated by insulators, inserted into the zebrafish genome using PhiC31 recombination at a single, known, landing pad site. In this line, NRE can activate expression of eGFP, while HS5 can activate mCherry (Figure 1C). Thus, the activity of these enhancers during zebrafish development can be visualised by the expression of eGFP and mCherry.

We used time-lapse and live imaging of these NRE-eGFP/HS5-mCherry enhancer-reporter embryos to compare the spatiotemporal activities of HS5 and NRE during development. This recapitulates the known domains of activity for these elements. At 24 hours post fertilisation (hpf) NRE-eGFP is active within the retina, while HS5-mCherry can be visualised in the retina as well as the neural tube and forebrain (Figure 1C, Supplementary movie 1). The activity of both of these elements in the retina peaks between 24-48 hpf, and decreases thereafter (Supplementary movie 1). We confirmed this result by creating a dye-swapped dual enhancer-reporter line using random Tol2 integration on a wild type background, in which NRE now activates mCherry while HS5 can activate eGFP (NRE-mCherry/HS5-eGFP) (Figure 1D).

In both reporter lines, within the retina, we observed an apparent enrichment of HS5 activity specifically in the temporal portion of the retina (Figure 1C-E). This was quantified by comparison of the mean eGFP and mCherry fluorescence in the two sides of the retina. There was no significant difference in overall fluorescence intensity for NRE between the nasal and temporal sides of the retina, but HS5 activity was significantly higher in the temporal part of the retina (Figure 1F).

To investigate this further, we carried out high-resolution live imaging of multiple NRE-eGFP/HS5-mCherry enhancer-reporter embryos at 24, 48, and 72 hpf. We observed a clear difference in the spatial activity of these enhancers within the retina, with NRE broadly active throughout and HS5 highly active in the temporal portion at all three time-points (Figure 2A). Quantification of mean fluorescence intensity confirmed significantly higher HS5 (mCherry) signal in the temporal versus nasal retina from 24-72 hpf, which was not observed for NRE (eGFP) (Figure 2B).

**Figure 2.**
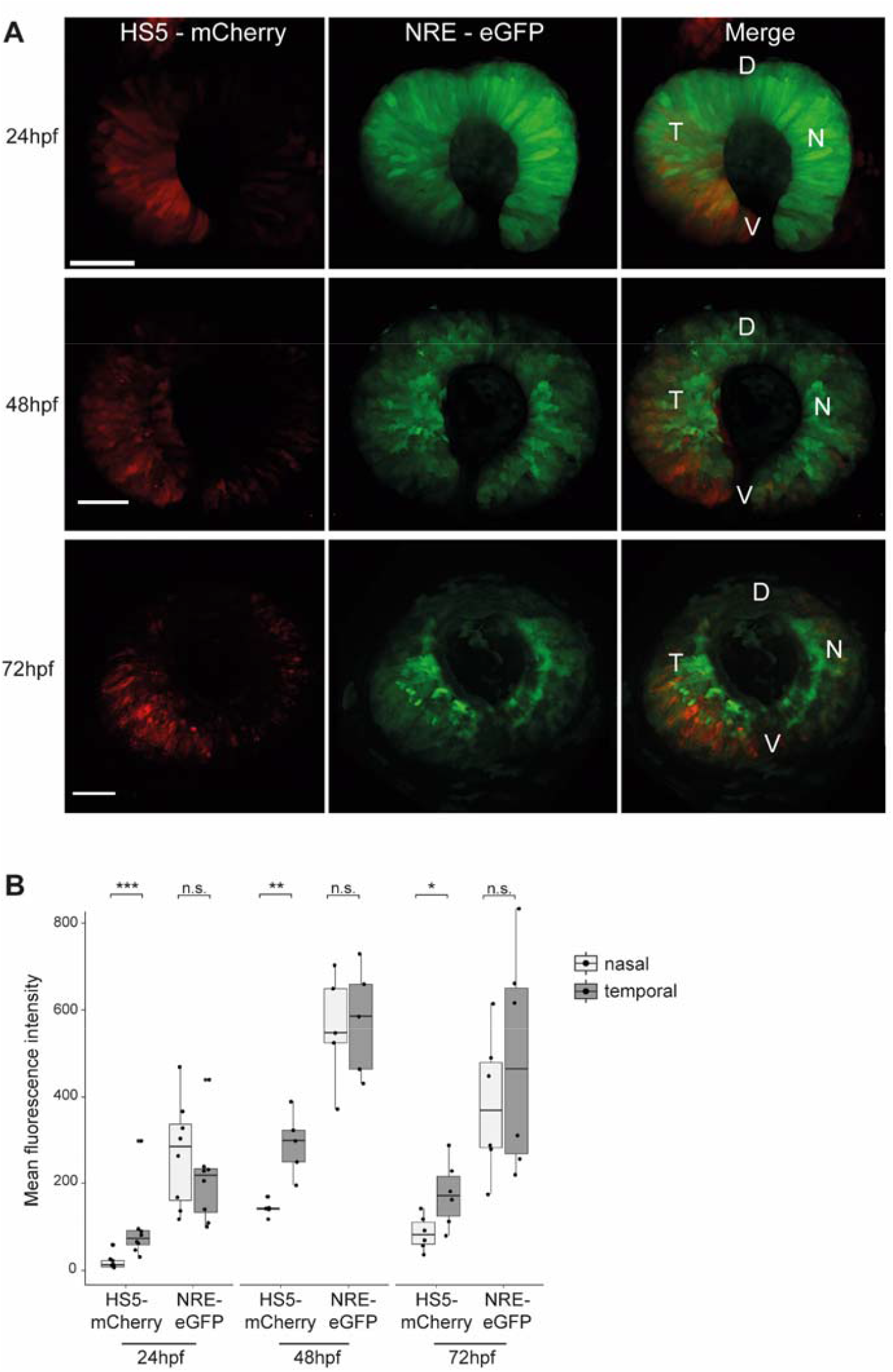
HS5 and NRE are active in different zones of the developing retina. **(A)** Live imaging at 24, 48, and 72 hpf in the developing retina of NRE-eGFP/HS5-mCherry F1 embryos. D, dorsal; V, ventral; T, temporal; N, nasal. Scale bars 50 µm. **(B)** Quantification of mean fluorescence intensity for mCherry (HS5) and eGFP (NRE) in the nasal versus temporal retina at 24, 48, and 72 hpf. Each measurement represents one embryo. The activity of HS5 (mCherry signal) is significantly higher in the temporal retina at all time points. n F1 embryos imaged ≥ 6 for all time points. Wilcoxon test results: ns, not significant; *, p<0.05; **, p<0.01; ***, p<0.001.

As a control, we created a dual enhancer-reporter line containing the NRE enhancer at both positions in the cassette (NRE-eGFP/NRE-mCherry) (Supplementary figure 1A). Imaging of these embryos showed a clear overlap between eGFP and mCherry expression throughout the retina (Supplementary figure 1A). Quantification of mean fluorescence intensity at 48 and 72 hpf showed no significant difference in the temporal versus nasal signal for mCherry or eGFP (Supplementary figure 1B). However, we observed that the fluorescence intensity for NRE-mCherry at 24 hpf was significantly higher in the nasal retina (Supplementary figure 1B). This is consistent with the NRE-eGFP and NRE-mCherry measurements in the HS5/NRE reporter lines, which show a modest but not significant increase in NRE activity in the nasal versus temporal retina at 24 hpf (Figures 1F and 2B). We conclude that at 24 hpf NRE activity is modestly higher in the nasal retina, the direct inverse of HS5 whose activity is highest in the temporal retina from 24-72 hpf. These contrasting patterns of activity led us to speculate that HS5 and NRE have distinct functions in *PAX6*-mediated retinal development.

### Single-cell RNA sequencing reveals that HS5 and NRE are active in distinct cell types in the developing zebrafish retina

In order to define the precise cell types within the retina where HS5 and NRE are active, we carried out scRNA-seq on retinae from NRE-eGFP/HS5-mCherry reporter embryos. With this technique, we aimed to uncover cell type or transcriptional differences between the two enhancer-active populations. We dissected eyes from 48 hpf NRE-eGFP/HS5-mCherry embryos (F1) and used FACS to enrich for either mCherry-positive/HS5-active cells or eGFP-positive/NRE-active cells. Three samples for each population were processed for scRNA-seq using the 10x Genomics Chromium single cell 3’ gene expression technology. After quality control (QC) and filtering we carried out K-nearest neighbour analysis followed by Louvain clustering on 6,288 cells, revealing 13 distinct clusters (Figure 3A). We used enriched marker genes to annotate each cluster, using gene expression data from ZFIN and the literature for known retinal cell types (Sprague et al., 2008) (Figure 3B-C, Supplementary figure 2, Supplementary table 1). A large proportion of the cells were classified as progenitors or stem cells, marked by high expression of *pcna* (proliferating cell nuclear antigen). As expected, these clusters were also characterised as actively proliferating (S and G2/M), in contrast to the more differentiated cell type clusters (Supplementary figure 3A). The stem cell cluster has high expression of genes such as *fbl* and *fabp11a* (Watanabe-Susaki et al., 2014), while the general retinal progenitor cluster is marked by high expression of the Notch signalling genes *notch1a* and *hes2*.*2*, and the progenitor marker *foxn4* (Li et al., 2004). In addition to these cell types, we identified clusters of amacrine, horizontal, bipolar, Müller glia, photoreceptor (PR), and retinal ganglion cells (RGCs). We were also able to annotate sub-types such as a cluster of OFF bipolar cells expressing *fezf2* (Suzuki-Kerr et al., 2018), and cholinergic amacrine cells, distinguished from the general amacrine cell cluster by expression of *sox2* (Whitney et al., 2014) (Figure 3A,B).

**Figure 3.**
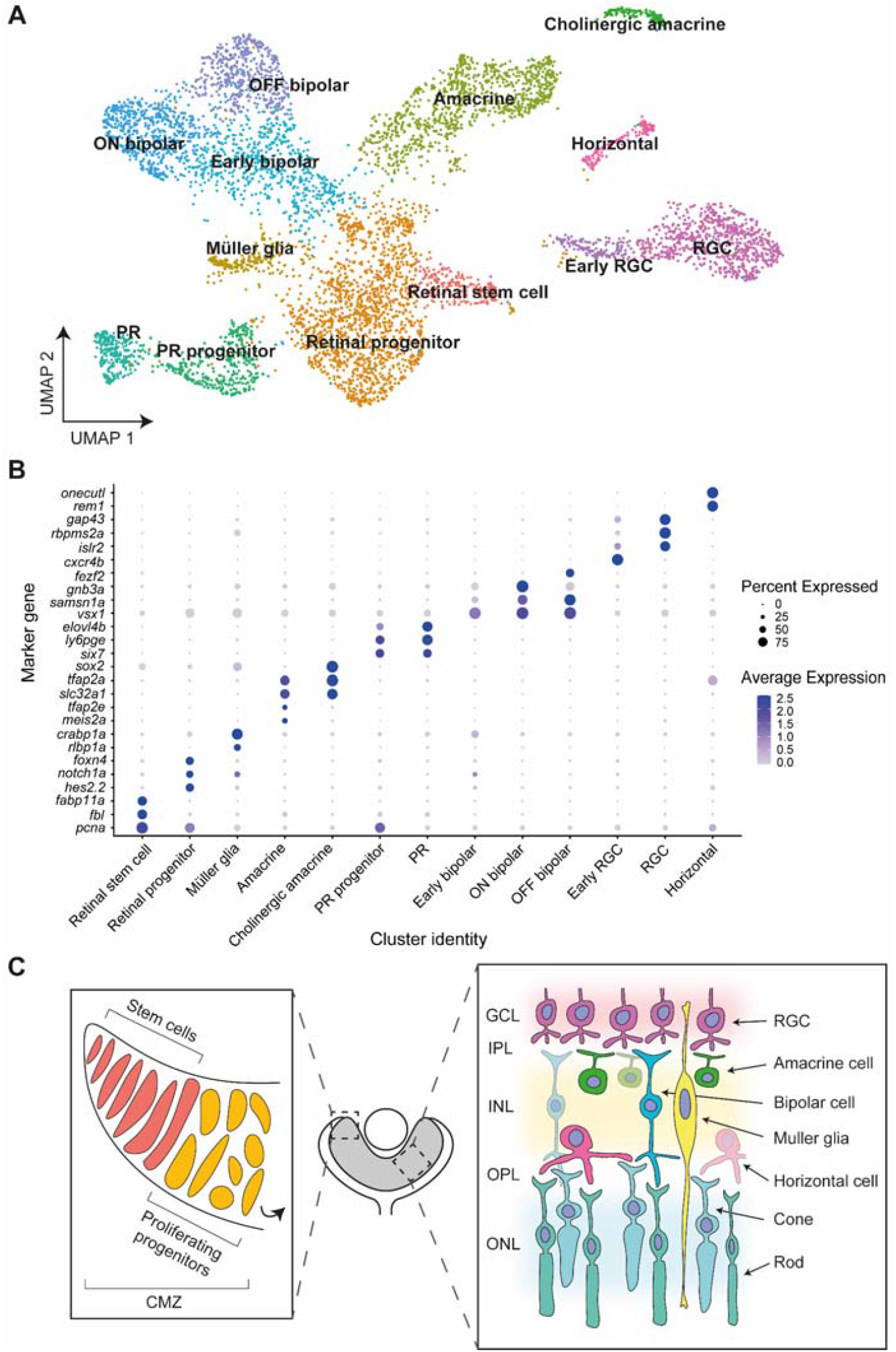
Single-cell RNA sequencing of NRE-eGFP/HS5-mCherry retinal cells at 48 hpf. **(A)** Cells from 6 sequenced libraries of NRE-eGFP/HS5-mCherry F1 embryos at 48 hpf, merged into one sample, and visualised on a Uniform Manifold Approximation and Projection (UMAP) plot created by Louvain clustering using Seurat (Butler et al., 2018). Retinal cell types are manually annotated to clusters depending on marker gene expression. **(B)** Dot plot showing average expression level and percentage of cells expressing key marker genes used to annotate cell type clusters. **(C)** Schematic of cell types in the zebrafish retina. In the ciliary marginal zone (CMZ) retinal stem cells undergo asymmetrical division to give rise to a rapidly proliferating pool of retinal progenitor cells (left), which divide and differentiate to form cells present in the retinal layers (right) (Richardson et al., 2016; Wan et al., 2016). Colours correspond to the annotated cluster cell types in **(A)**. RGC, retinal ganglion cell; GCL, ganglion cell layer; IPL, inner plexiform layer; INL, inner nuclear layer; OPL, outer plexiform layer; ONL, outer nuclear layer.

We also used the zebrafish single-cell transcriptomic atlas to assign cell type identities in our dataset (Farnsworth et al., 2019). The majority of cells were assigned to ‘retinal differentiating’ and ‘retinal progenitor’ identities (Supplementary figure 3B). The ‘retinal progenitor’ identity appears to specifically mark cells from the retinal stem cell cluster (Supplementary figure 3B). Comparing the assigned atlas identities to our cell type annotations in more detail proved informative. For example, the atlas identity RetNeuron25 appears to be specific to amacrine cells, while RetDiff25e appears to mark RGCs (Supplementary figure 3C).

In order to confidently identify the cell types where the HS5 and NRE enhancers are active, we looked for the presence of mCherry and eGFP reads, respectively. We observed the highest expression of eGFP reads within both the amacrine and stem cell clusters, and the highest expression of mCherry reads within the Müller glia and retinal progenitor/stem cell clusters (Figure 4A-C). Importantly, all of these cell type clusters also have high expression of zebrafish *pax6a* and *pax6b*, making them conceivable candidates for *PAX6* enhancer-active cell types (Supplementary figure 4). Although we used FACS to enrich for eGFP and mCherry-positive cells, most cells within our dataset do not express eGFP or mCherry. We expect the reason for this to be two-fold. Firstly, ‘dropout’ of reads in scRNA-seq data is common, particularly if the gene is lowly expressed and there is low mRNA in individual cells, which we expect in our samples (Kharchenko et al., 2014). It is likely that ‘dropout’ of eGFP and mCherry reads has occurred in clusters where we see highest expression of these genes. Secondly, false-positive selection can occur during FACS, particularly where there is autofluorescence within the sample, which is true for zebrafish eyes (Shi et al., 2009).

**Figure 4.**
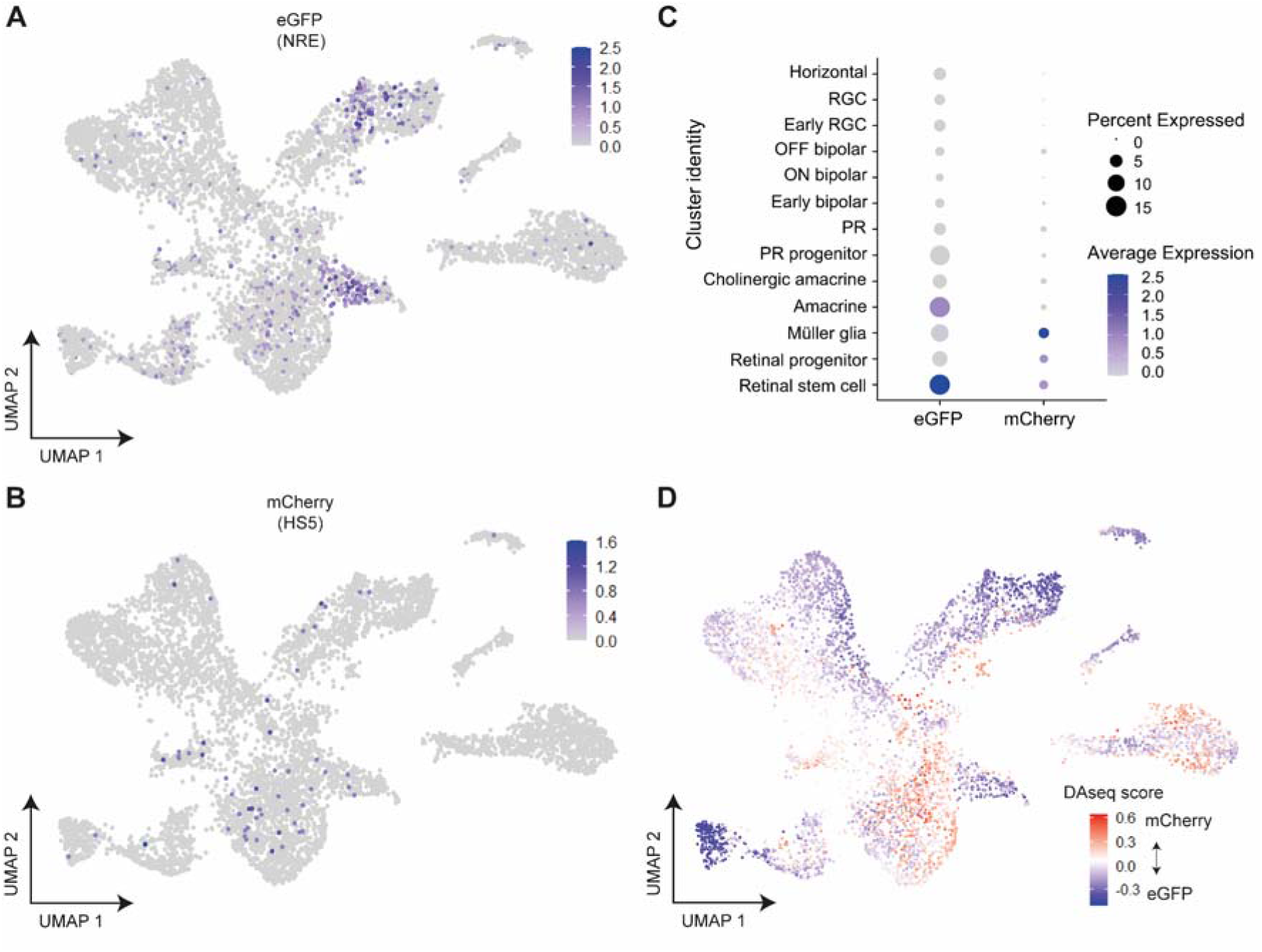
Assigning the identity of HS5 and NRE-active cells using the expression of eGFP and mCherry, and differential abundance analysis in cell type clusters. **(A)** Expression of eGFP and **(B)** mCherry in single cells visualised on UMAP plots (clustered as in Figure 3A). eGFP expression is enriched in amacrine and retinal stem cell clusters. **(C)** Dot plot showing average expression and percentage of cells expressing mCherry (HS5-active) and eGFP (NRE-active) in cell type clusters. Enrichment is seen for eGFP expression in amacrine and retinal stem cell clusters, and for mCherry in Müller glia and retinal progenitor/stem cell clusters. **(D)** DAseq (Zhao et al., 2021) differential abundance analysis comparing the relative prevalence of cells from mCherry-enriched or eGFP-enriched samples. Cells are coloured by DAseq score and displayed on a UMAP plot. A score is calculated for each cell based on the abundance of cells from both populations in the cell’s neighbourhood. Positive (red) scores indicate an abundance of cells from mCherry-enriched samples; negative (blue) scores indicate an abundance of cells from eGFP-enriched samples.

Because of the lower than expected number of eGFP and mCherry reads in our scRNA-seq dataset, we employed differential abundance analysis to validate the mCherry-enriched (HS5) and eGFP-enriched (NRE) clusters. We used two statistical methods of differential abundance analysis (DAseq and MiloR) to calculate the relative abundance of cells from eGFP or mCherry-enriched samples within regions of our Seurat clusters (Dann et al., 2022; Zhao et al., 2021). Both of these methods showed regions of enrichment for cells from mCherry-enriched samples within retinal progenitor and Müller glia clusters, and enrichment for cells from eGFP-enriched samples within the amacrine and stem cell clusters when visualised on a UMAP plot (Figure 4D and Supplementary figure 5A), and in the DAseq scores per cluster (Supplementary figure 5B).

We also applied topic modelling (Dey et al., 2017) to our dataset to simultaneously identify gene topics (corresponding to sets of coexpressed genes) and cell-topic weights (which quantify the proportion of a cell’s transcriptome described by a given gene topic). We then used a two-sided Pearson’s product-moment correlation test to find topics correlated with expression of mCherry and eGFP (Supplementary table 2). We find that topic 1 from this analysis was most highly correlated with mCherry expression, and contributes highly to cells from Müller glia and progenitor clusters (Supplementary figure 5C). Inspecting the genes of this topic reveals that it is enriched for notch signalling genes and Müller glia markers, again giving confidence that mCherry-expression/HS5-activity appears enriched in cells of the progenitor and Müller glia clusters (Supplementary table 3). The topic most highly correlated with expression of eGFP (topic 5) contributes highly to cells in the amacrine clusters, and again is enriched for genes characteristic of amacrine cells, including *tfap2a* and *slc32a1* (Supplementary figure 5D, Supplementary table 4). Thus, the topics most highly correlated with eGFP and mCherry expression correspond to topics that are characteristic of the cell type clusters we find using differential abundance analysis. The agreement between the cluster-specific enrichment of eGFP and mCherry reads with the differential abundance analysis and topic modelling, suggests that our coupling of the QSTARZ assay with scRNA-seq analysis is a powerful way of identifying the precise cell types within a tissue where enhancers are active.

### HS5 and NRE-active cell types are confirmed by immunofluorescence

Our scRNA-seq dataset indicates that NRE is primarily active in amacrine and retinal stem cells, while HS5 appears to be active in proliferating progenitors and Müller glia (Figure 4). This is in agreement with our live imaging data detecting NRE-eGFP signal in the distal tip of the ciliary marginal zone (CMZ) and in cells of the inner nuclear layer (INL), and HS5-mCherry signal in the more proximal region of the CMZ and in cells of the INL, at 48 hpf (Figure 5A). To confirm the identity of these enhancer-active cell types we carried out immunofluorescence on eye sections from NRE-eGFP/HS5-mCherry embryos. We stained for PCNA – a marker of progenitor and stem cells in the CMZ, HuC/D (encoded by *elavl3/4*) – a marker for RGCs in the ganglion cell layer (GCL) and amacrine cells in the INL, and glutamine synthetase (GS, encoded by *glula*) – a marker for Müller glia (Ekström and Johansson, 2003; Thummel et al., 2008; Fischer et al., 2013). Our scRNA-seq data confirms the expression of these genes within these specific cell types in our samples (Figure 5B).

**Figure 5.**
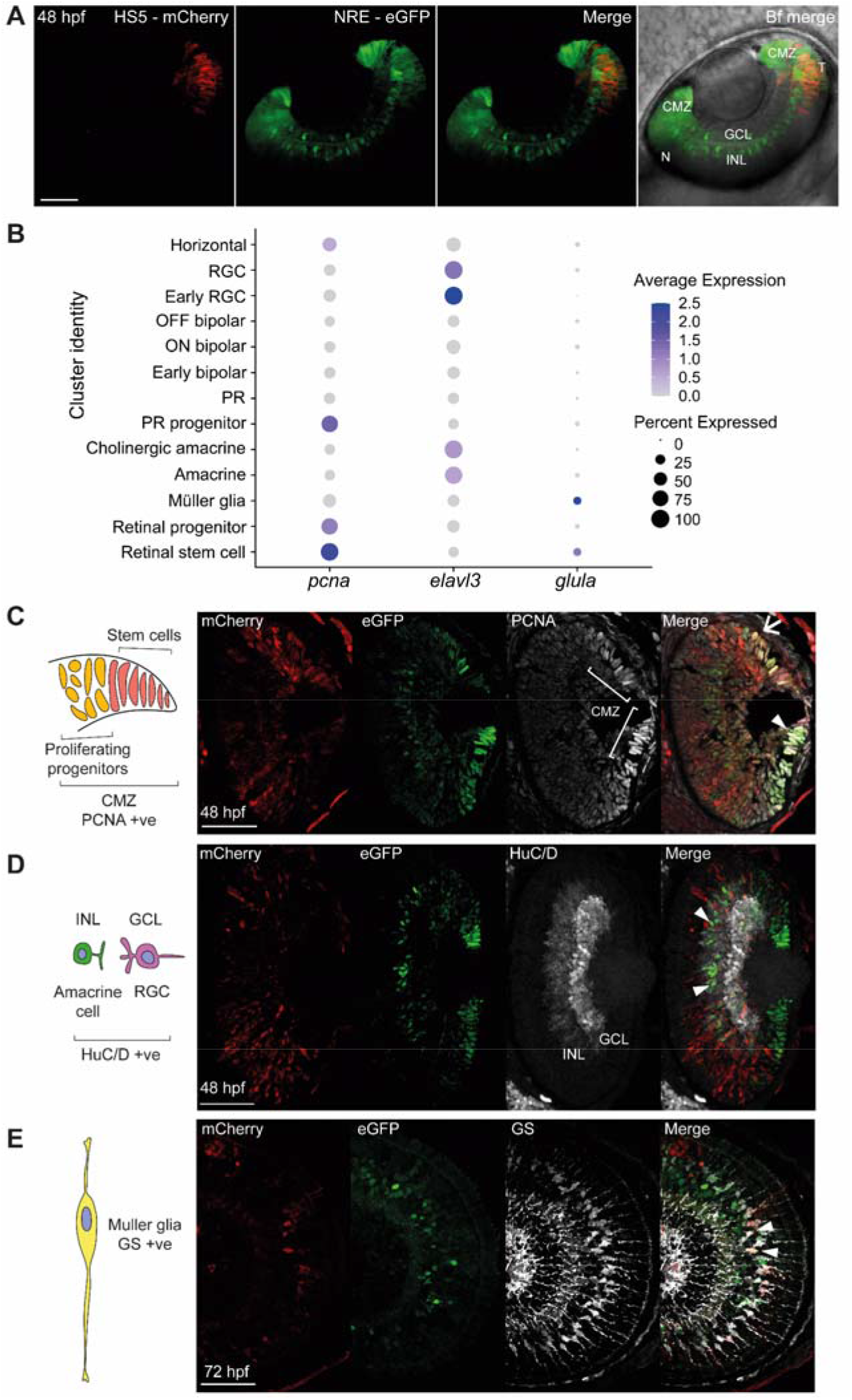
Immunofluorescence identifies enhancer-active cell types. **(A)** Coronal-orientation image of a NRE-eGFP/HS5-mCherry embryo at 48 hpf showing the activity of NRE (eGFP) in the distal CMZ (stem cell niche) and cells of the INL, and HS5 (mCherry) activity in the temporal retina, in the proximal CMZ and cells of the INL. **(B)** Dot plot showing average expression and percentage of cells expressing *pcna, elavl3* (encoding HuC), and *glula* (encoding glutamine synthetase (GS)) in cell type clusters. **(C)** Immunofluorescence for PCNA, mCherry, and eGFP on a coronal eye section from an NRE-eGFP/HS5-mCherry F1 embryo at 48 hpf. PCNA is a marker for progenitors and stem cells in the CMZ. Arrow indicates an mCherry/PCNA positive cell. Arrowhead indicates a eGFP/PCNA positive cell. **(D)** As in **(C)** but using an antibody detecting HuC/D (*elavl3/4*). HuC/D is a marker for RGCs in the GCL and amacrine cells in the INL. Arrowheads indicate eGFP/HuC/D positive cells. **(E)** As in **(C)** but using an antibody detecting glutamine synthetase (GS), on an embryo at 72 hpf (sagittal section). GS is a marker for Müller glia. Arrowheads indicate mCherry/GS positive cells. Scale bars 50 µm.

We detect strong expression of eGFP in PCNA+ cells of the CMZ, particularly in the stem cell region in the distal tip (Figure 5C, arrowhead). We also observed mCherry expression in PCNA+ cells, however in contrast to eGFP this is seen in the more proximal region of the CMZ, where the rapidly proliferating progenitors are located (Figure 5C, arrow). Staining for HuC/D, we observed co-staining with eGFP in amacrine cells of the INL (Figure 5D, arrowheads). We also detected mCherry-expression in a subset of GS+ Müller glia at 72 hpf (Figure 5E, arrowheads). These results confirm our findings that HS5 and NRE are active in distinct cell types of the developing retina, with NRE active in amacrine and retinal stem cells, and HS5 active in proliferating progenitors and Müller glia.

## Discussion

Several issues challenge our understanding of the precise role of enhancers in developmental and disease processes, including relating the sequence-specific grammar of these elements to their function, defining their exact domains of action, and disentangling the additive, redundant or distinct functions of apparently similar enhancers. A few enhancers, such as the famous ZRS limb enhancer of *SHH*, have been so thoroughly characterised that they can now be manipulated at the single base pair level to achieve specific phenotypes (Lettice et al., 2003; Kvon et al., 2020; Lim et al., 2022). However, our understanding of the other hundreds of thousands of predicted enhancers in the human genome lags far behind this, and for the majority it remains challenging to identify even the target gene with confidence. Effective and efficient in vivo analysis pipelines for characterisation of regulatory elements are needed to address this knowledge gap. In this study we have described an in vivo method for generating high-resolution data on the precise activities of developmental enhancers. Using a combination of live imaging and scRNA-seq in embryos derived from dual enhancer-reporter transgenic zebrafish lines, we clearly demonstrate distinct patterns of spatiotemporal and cell type specific activity during retinal development for two retinal enhancers (HS5 and NRE) from the *PAX6* regulatory region (Figure 6).

**Figure 6.**
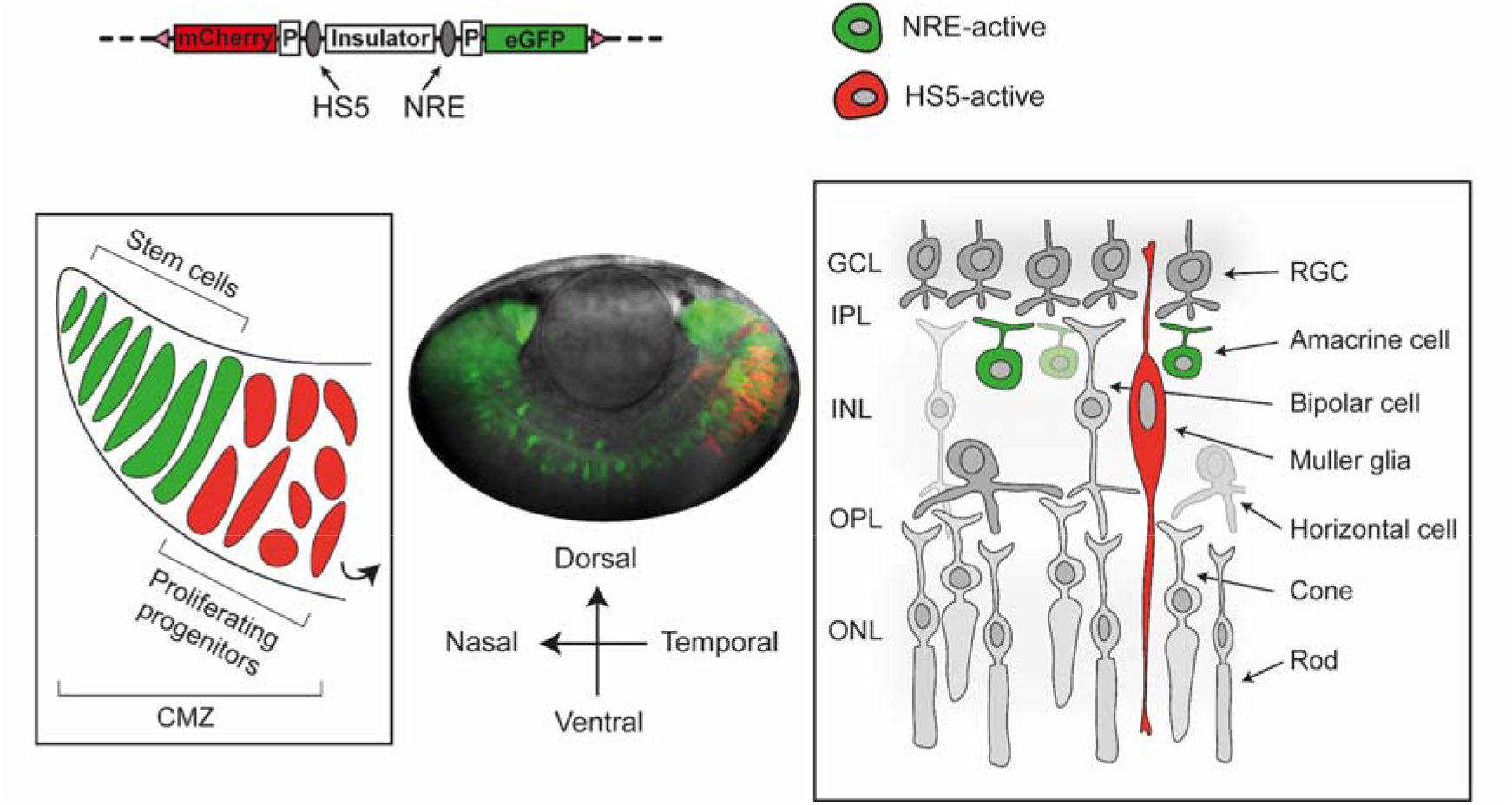
Investigating the distinct functions of overlapping *PAX6* retinal enhancers in a zebrafish dual enhancer-reporter assay. *PAX6* retinal enhancers HS5 and NRE have distinct spatiotemporal and cell type specific functions in a dual enhancer-reporter system in zebrafish embryonic development. NRE is active throughout the developing retina, and is localised to stem cells of the CMZ and amacrine cells of the INL (green). HS5 is active mainly in the temporal region of the retina, in proliferating progenitors and Müller glia (red).

It has been shown that NRE is active in a retinal progenitor population during mouse embryonic development (Marquardt et al., 2001). In this study, we have shown that NRE is specifically active in retinal stem cells of the CMZ, as can be seen by scRNA-seq (Figures 3 and 4), and high-resolution live-imaging and immunofluorescence (Figure 5). NRE has also been reported as active in non-cholinergic amacrine cells in postnatal mouse eyes, and necessary for their development (Kim et al., 2017). This mirrors the known functions of PAX6, which is essential for the generation of amacrine cells (Remez et al., 2017). Here we show that as early as 48 hpf in the zebrafish (∼E14.5 in mouse), NRE is seen to be strongly active in differentiating amacrine cells by immunofluorescence imaging (Figure 5), in specifically the non-cholinergic population, as is shown by scRNA-seq (Figures 3 and 4). It is likely that the reduced amacrine cell phenotype reported by Kim et al. (2017) upon NRE deletion, is due to the loss of NRE activity at these early time points affecting *PAX6* expression. The dual activity of NRE in retinal stem cells and differentiating amacrine cells reflects the multifarious functions of *PAX6*, which is necessary both for promoting the proliferation and potency of retinal progenitors, as well as directing cell-cycle exit and differentiation of cell types including amacrine cells (Farhy et al., 2013; Remez et al., 2017).

Previously, little was known about the retinal function of HS5 (McBride et al., 2011). Here we show that HS5 is active in the rapidly proliferating progenitor population of the developing retina, as well as Müller glia (Figures 3, 4 and 5). Transcriptionally, Müller glia share similarities with retinal progenitor cells, in that they are specialised glial cells with progenitor potential (Jadhav et al., 2009). Again, it is known that *PAX6* expression is detected in both of these cell types, and is necessary for the maintenance of progenitors and generation of Müller glia (Marquardt et al., 2001; Joly et al., 2011). In zebrafish, Müller glia are capable of undergoing transcriptional reprogramming to produce retinal progenitor cells after acute injury to the retina (Goldman, 2014). This property has also been shown for human Müller glia in vitro and in rodent transplants (Singhal et al., 2012). During development, each retinal progenitor cell is capable of forming several neural retina cells and a single Müller glia (Rulands et al., 2018). Whether HS5 is active in Müller glia at later stages, or if the activity of HS5 in these cells is linked to Müller glia development and HS5 activity in upstream retinal progenitors is unclear.

HS5 and NRE also showed distinct differences in their spatiotemporal patterns of activity. The activity of HS5 in the temporal retina is notable, as in zebrafish this region is a fovea-like region of high-acuity, characterised by specialisation and increased density of cell types (Schmitt and Dowling, 1999; Yoshimatsu et al., 2020). It is unclear if HS5 may also show a similar pattern of activity in the human fovea, and whether this may be functionally relevant to fovea development. A fovea-like region in mice was only recently identified and requires further characterisation (van Beest et al., 2021). The previous study identifying HS5 in a mouse lacZ reporter assay did not report any specific spatial activities for this enhancer within the developing eye, and the resolution of this assay would be poorly suited to do this (McBride et al., 2011).

A limitation of this study, and enhancer-reporter assays in general, is the fact that the enhancers are tested outside of their native genomic context. An advantage of this dual enhancer-reporter assay in zebrafish however is that the developmental context of enhancer activity is preserved, and this can be easily followed in live embryos, particularly in developing eyes. As such, zebrafish are an ideal model to study the activity of the human *PAX6* retinal enhancers tested here. Zebrafish are already a well-established model for studying ocular genetics, as eyes and eye development are highly conserved between zebrafish and humans, including morphology, cell-types, protein-markers and gene expression (Richardson et al., 2016; Angueyra and Kindt, 2018). Species specific differences may limit the interpretation of some results, which we have discussed.

To the best of our knowledge, ours is the first report where the activities of developmental enhancers have been mapped in vivo at single-cell resolution to reveal distinct patterns of activity. We have shown that two overlapping *PAX6* retinal enhancers have distinct spatial, temporal, and cell type specific activities, and as such it is highly likely that they are non-redundant with differing functions in human retinal development. Based on our results, we suggest that other enhancers with apparently overlapping domains of activity based on lower resolution assays, may have similar distinct functions that require proper dissection at high spatial and temporal resolution. Indeed, there are several other retinal enhancers at the *PAX6* locus that have yet to be characterised, and which may also have unique or redundant functions. The comprehensive framework described here to evaluate HS5 and NRE provides a systematic pipeline to investigate the function of enhancers, and the consequences of enhancer mutations. The scRNA-seq dataset of developing zebrafish retinal cell types, in which different enhancers are active, sheds light on gene regulatory networks during retinal development, and can be utilised as a valuable resource in future studies.

## Materials and methods

### Generation of dual enhancer-reporter constructs

Dual enhancer-reporter constructs were generated using Gateway cloning (Invitrogen) as described in Bhatia et al. (2021). Destination vectors containing the gata2-eGFP and gata2-mCherry gene units were synthesised by GeneArt. These contain R3/R4 Gateway recombination sites for insertion of enhancer sequences, and the cassette is flanked by either PhiC31 or Tol2 recombination sites for genome integration. The sequences of HS5 and NRE were PCR amplified from human genomic DNA using Phusion high fidelity polymerase (NEB). Hg38 genome coordinates and sequences of primers used, containing overhang Gateway recombination sequences, are in Supplementary table 4. Purified PCR products were cloned into Gateway pDONR entry vectors (pP4P1r or pP2rP3) using BP clonase. Plasmids were sequenced using the original enhancer primers to verify integration. The insulator construct was previously created by cloning into a pDONR221 vector (Bhatia et al., 2021). The insulator construct used in this study contains 2.5 copies of the chicken HS4 sequence. Final dual enhancer-reporter constructs were created using LR clonase in a multi-way Gateway reaction, to combine the two enhancer sequences and the insulator into the destination vector. The NRE-eGFP/HS5-mCherry and NRE-eGFP/NRE-mCherry constructs were created using the PhiC31 destination vector, and the swap construct NRE-mCherry/HS5-eGFP was created using the Tol2 destination vector.

### Generation of transgenic zebrafish lines

Transgenic zebrafish lines were created as described in Bhatia et al. (2021). For the PhiC31 constructs, embryos were obtained from landing-line adults and injected at the one-cell stage. For the Tol2 construct, embryos were obtained from wild type adults (strain AB) and injected at the one-cell stage. F0s were screened for mosaic expression of eGFP and mCherry (and loss of landing-line fluorescence expression when using PhiC31), and raised to adulthood. Sexually mature F0s were crossed with wild type adults and F1s screened for fluorescence using a Leica M165FC fluorescence stereo microscope. Fluorescent F1s were used for imaging and scRNA-seq. At least two F0 founders per construct were used to generate F1s.

### Zebrafish husbandry

Adult zebrafish were maintained according to standard protocols (Sprague et al., 2008). Embryos were raised at 28.5°C and staged by hpf and morphological criteria (Kimmel et al., 1995). All zebrafish work was carried out under a UK Home Office licence under the Animals (Scientific Procedures) Act 1986.

### Live and time-lapse imaging

Prior to imaging, all embryos were treated with 0.003% 1-phenyl2-thio-urea (PTU) from 12 hpf to prevent pigmentation from developing. For live imaging, embryos at the correct stage were anaesthetised with Tricaine (20–30 mg/l) and mounted in 1% low-melting point (LMP) agarose in a glass-bottom dish (Cellvis P06-1.5H-N). Embryo media was added to the dish to prevent drying out. Embryos were imaged using a Nikon A1R (scanning) confocal microscope, using a 10x objective or 40x water immersion objective. Data were acquired using NIS Elements AR software (Nikon Instruments Europe). Images were taken as a Z stack (1 µm step size). Imaging was repeated for several F1s per line (see figure legends). For time-lapse imaging, embryos were mounted as described, and a portion of the LMP agarose surrounding the embryo head/body was carefully cut away using a microsurgical knife (World Precision Instruments). This left only the tail of the embryo embedded in agarose, allowing unimpeded development throughout the imaging time-course. Tricaine (20–30 mg/l) and PTU (0.003%) were added to the embryo media covering the embryos. Time-lapse imaging was carried out using an Andor Dragonfly (spinning disk) confocal (Andor technologies), using a 10x objective. Data were collected in Spinning Disk 40µm pinhole mode on the iXon 888 EMCCD camera using Andor Fusion acquisition software. Embryos were maintained at 28.5°C using an Okolab bold line stage top incubation chamber (Okolab S.R.L). Images were taken as a Z stack (1 µm step size) every 60 minutes for ∼ 48 hours.

### Immunostaining and imaging

NRE-eGFP/HS5-mCherry embryos were dechorionated and fixed in 4% paraformaldehyde (PFA) overnight at 4°C, then stored in 30% sucrose. Embryos were embedded in optimal cutting temperature compound (OCT), and flash-frozen at -80°C. OCT blocks were cryosectioned and samples dried onto SuperFrost Plus Adhesion slides (Epredia). Samples were then rehydrated in phosphate-buffered saline (PBS) for 5 minutes at room temperature (RT), followed by antigen retrieval in 10 mM sodium citrate (pH 6), heated for 20 minutes in a rice cooker. Once returned to RT, samples were washed 3 time in PBS 0.1% Triton X-100 (PBST) for 5 minutes at RT. Samples were blocked for 1 hour at RT in 1% bovine serum albumin (BSA), 10% goat serum in PBST, then incubated with primary antibody in blocking solution overnight at 4°C. All samples were incubated with chicken anti-GFP (Gene Tex, GTX13970) (1:1000), rabbit anti-mCherry (Proteintech, 26765-1-AP) (1:1000), and either mouse anti-PCNA (Sigma, P8825) (1:200), mouse anti-HuC/D (Invitrogen, A21271) (1:200), or mouse anti-GS (Proteintech, 66323-1-Ig) (1:300). After incubation slides were washed 3 times in PBS for 20 minutes at RT, followed by incubation with secondary antibody for 2 hours at RT. The following Alexa Fluor-conjugated secondary antibodies were used at 1:1000 dilution – goat anti-mouse Alexa Fluor 647 (Invitrogen, A-21235), goat anti-chicken Alexa Fluor 488 (Invitrogen, A-11039), goat anti-rabbit Alexa Fluor 546 (Invitrogen, A-11035). Samples were then washed 3 times in PBS for 20 minutes at RT, with 1 μg/ml of 4′,6-diamidino-2-phenylindole (DAPI) added to the last wash. Slides were mounted with a coverslip using mounting medium (Abcam, Ab104139), then stored in the dark at 4°C. Slides were imaged on a Zeiss LSM 900 scanning confocal using a 40x water immersion objective, in z-stack acquisition mode (0.5 μm step size).

### Image analysis

All images were processed using FIJI and are displayed in figures as maximum intensity projections (Schindelin et al., 2012). eGFP and mCherry mean fluorescence intensity measurements were taken using FIJI in the temporal and nasal portions of the retina only. In R, ggplot2 and ggpubr packages were used to plot these measurements and compare means between groups using a Wilcoxon test (Wickham, 2009).

### Dissociation and sorting of zebrafish embryonic retinae

F1 NRE-eGFP/HS5-mCherry embryos, and wild type embryos, were collected and treated with PTU as described. At 48 hpf, embryos were anaesthetised with Tricaine (20–30 mg/l) and placed into Danieau’s solution (Sprague et al., 2008). Eyes were dissected from ∼100-150 embryos using fine forceps (Dumont #5SF), and immediately placed into Danieau’s solution on ice. Samples were centrifuged at 300g for 1 minute at 4°C, then washed with Danieau’s solution. Washing step was carried out three times with Danieau’s solution, and once with FACSmax (Amsbio). In a final 500 µl FACSmax, the samples were passed through a 35 µm cell strainer to obtain single cell suspension (on ice). Samples were sorted for mCherry and eGFP fluorescence using a FACS Aria II (BD) or CytoFLEX SRT (Beckman Coulter) machine. Forward and side scatter sorting was used to select single cells from clumps and debris, and DAPI staining was used to exclude dead cells. A wildtype sample was used as a negative control for eGFP and mCherry fluorescence, to set the gates for sorting. Cells single positive for eGFP were selected for the eGFP samples. Due to the smaller population of mCherry positive cells, the yield of mCherry samples was increased by sorting cells single positive for mCherry or double positive.

### Single-cell RNA sequencing

After FACS, ∼10,000 cells/sample were processed using the 10x Genomics Chromium single cell 3’ gene expression technology (v3.1), according to the manufacturer’s instructions (Zheng et al., 2017). Sequencing was performed on the NextSeq 2000 platform (Illumina Inc.) using the NextSeq 1000/2000 P3 Reagents (100 cycles) v3 Kit. Sequencing data was processed using Cell Ranger (10x Genomics, v6.1.2). Cellranger mkfastq was used to create FASTQ files from raw sequencing data, followed by cellranger count to perform alignment, filtering, barcode counting, and UMI counting. A custom reference genome was created for alignment using cellranger mkref, combining the *Danio rerio* GRCz11 genome assembly with manually annotated eGFP and mCherry sequences.

### Computational analysis of scRNA-seq data

#### Cell calling and QC

Taking the raw (unfiltered) output from cellranger count, we used the emptyDrops function from DropletUtils to filter out empty droplets/barcodes not corresponding to cells (Lun et al., 2019). Mitochondrial and ribosomal genes were excluded from the emptyDrops analysis to improve the filtering of droplets containing ambient RNA or cell fragments. The scater package was used to filter cells based on the QC metrics of library size, detected genes, and mitochondrial reads (McCarthy et al., 2017). Cells with detected genes ≥ 500, library size ≥ 800, and mitochondrial reads ≤ 10% were retained.

#### Reference mapping and filtering

The SingleR package was used to annotate cell types based on mapping to the zebrafish single-cell transcriptome atlas (Aran et al., 2019; Farnsworth et al., 2019). Expression matrix and cell annotation data were downloaded from the UCSC cell browser (http://zebrafish-dev.cells.ucsc.edu); only the 2 dpf data were used for mapping. Erroneously sorted cells of non-retinal identity (for example pigmented cell types such as melanocytes with high autofluorescence) were filtered out at this stage. This was carried out to improve the resolution of clustering for retinal cell types.

#### Clustering and cell type annotation

Seurat (v4) was used for clustering and further analysis for a total of 6,288 cells (Butler et al., 2018). SCTransform was used to perform log-normalisation, scaling, and highly variable gene (HVG) detection on a dataset consisting of the 6 samples merged into one. Standard SCTransform options were used, with regression of mitochondrial expression and cell-cycle stage using ‘vars.to.regress’. We performed Principle Component Analysis (PCA) on the normalized counts matrix restricted to HVGs, using Seurat’s RunPCA function with number of PCs = 50. To enable integration of the samples, we then used Harmony to generate PCs corrected for batch effects between libraries (Korsunsky et al., 2019). The Harmony PCs were then used to perform K-nearest neighbour analysis (k=20) and Louvain clustering using Seurat (15 dimensions and resolution 0.6). Clusters were annotated as retinal cell types based on the highest expressed marker genes, and other known genes for each cell type, using information from the literature and ZFIN (Sprague et al., 2008). Cell cycle scoring was performed using the Seurat CellCycleScoring function, using zebrafish genes homologous to the ‘s.features’ and ‘g2m.features’ genes provided by Seurat.

#### Differential abundance analysis and topic modelling

DAseq and MiloR were used to perform differential abundance analysis between eGFP and mCherry-enriched samples using the standard, suggested parameters (Zhao et al., 2021; Dann et al., 2022). UMAP embeddings from Seurat were used as the graphing inputs. Topic modelling was carried out using fastTopics, using k=8 number of topics (Dey et al., 2017). Non-normalised counts were used as the input, as is standard for topic modelling. Correlation between fluorophore expression and topic scores were calculated using a two-sided Pearson’s product-moment correlation using the cor.test function from R stats (R Core Team, 2021).

## Acknowledgements

We thank A. Brombin and J. Travnickova for their guidance and advice on scRNA-seq, and helping with eye dissections alongside G. Alston, B. Bartlett, L. Gomez Acuna, A. Gonzalez Estevez and K. Purshouse. We thank E. Freyer and S. Campbell of the IGC FACS facility, and R. Clark of the Edinburgh Clinical Research Facility for processing 10x samples, and the MRC HGU Zebrafish facility and IGC Advanced Imaging Resource for support. Finally, we thank G. Alston and E. Friman for comments on the manuscript.

## Ethics

All zebrafish experiments were approved by the University of Edinburgh ethical committee and performed under UK Home Office license PPL PA3527EC3.

## Supplementary figures

**Supplementary figure 1.**
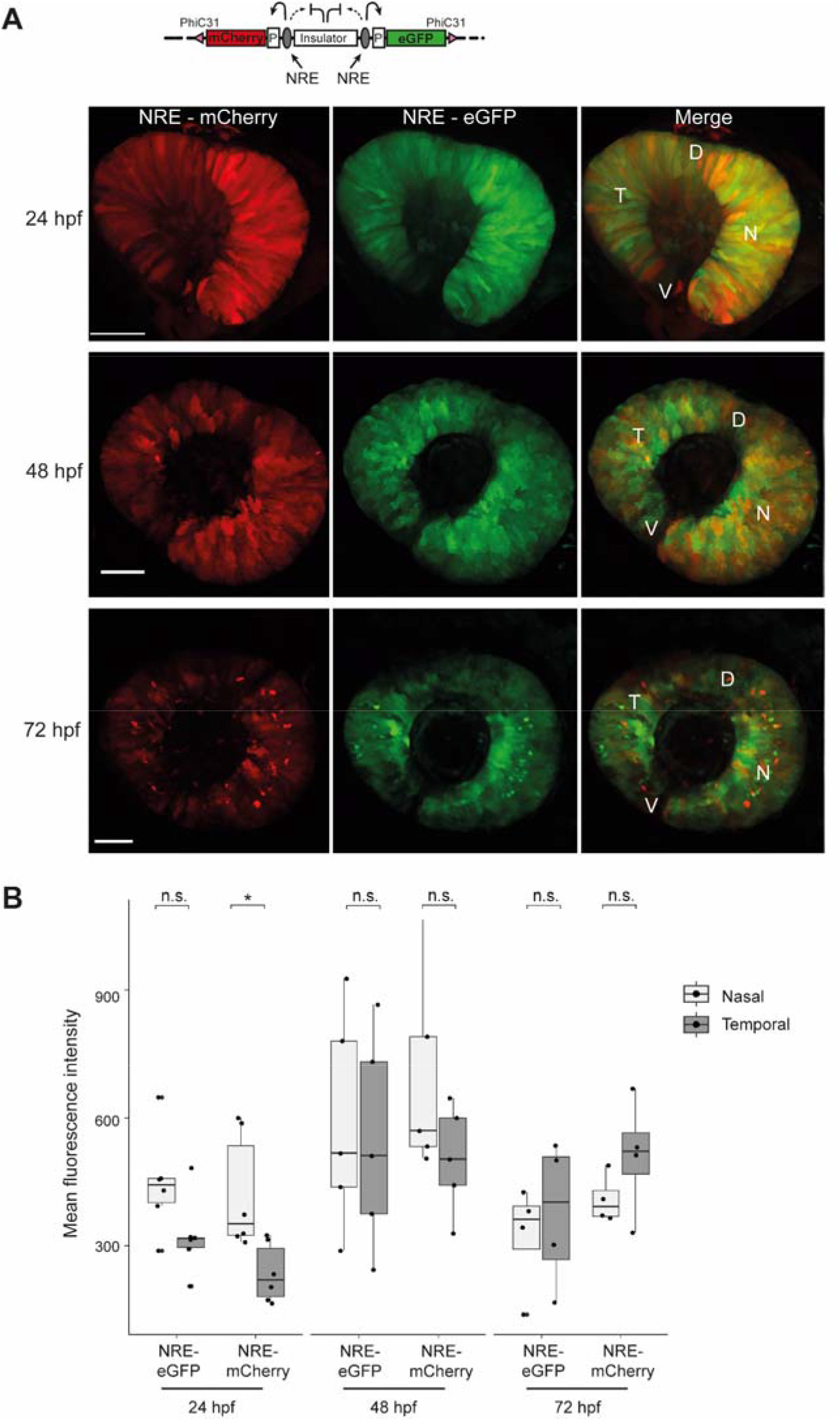
NRE control enhancer-reporter line reveals differential activity of NRE at 24 hpf. **(A)** Live imaging of NRE-eGFP/NRE-mCherry F1 embryos shows overlapping retinal signal for mCherry and eGFP at 24, 48, and 72 hpf. At 72 hpf a layer of strongly positive NRE-active cells can be seen in the neural retina. **(B)** Quantification of mean fluorescence intensity for mCherry and eGFP in NRE control embryos at 24, 48, and 72 hpf shows that NRE (mCherry) activity is significantly higher in the nasal retina compared with the temporal zone at 24 hpf. n F1 embryos imaged ≥ 5 for all time points. Wilcoxon test results: ns, not significant; *, p<0.05. D, dorsal; V, ventral; T, temporal; N, nasal. Scale bars 50 µm.

**Supplementary figure 2.**
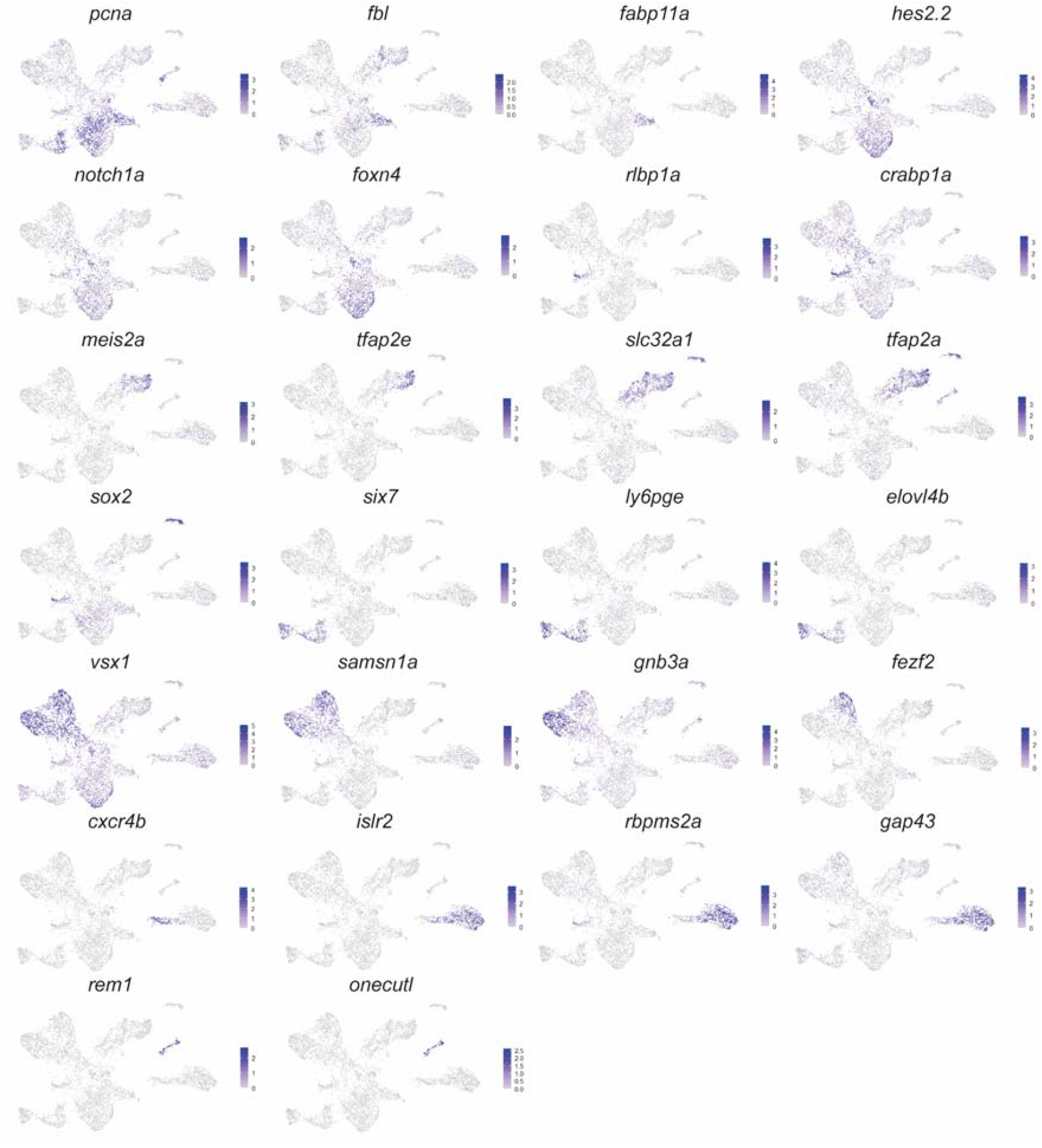
Expression of marker genes used for cluster annotation. Expression of cluster marker genes from Figure 3B visualised on UMAP plots.

**Supplementary figure 3.**
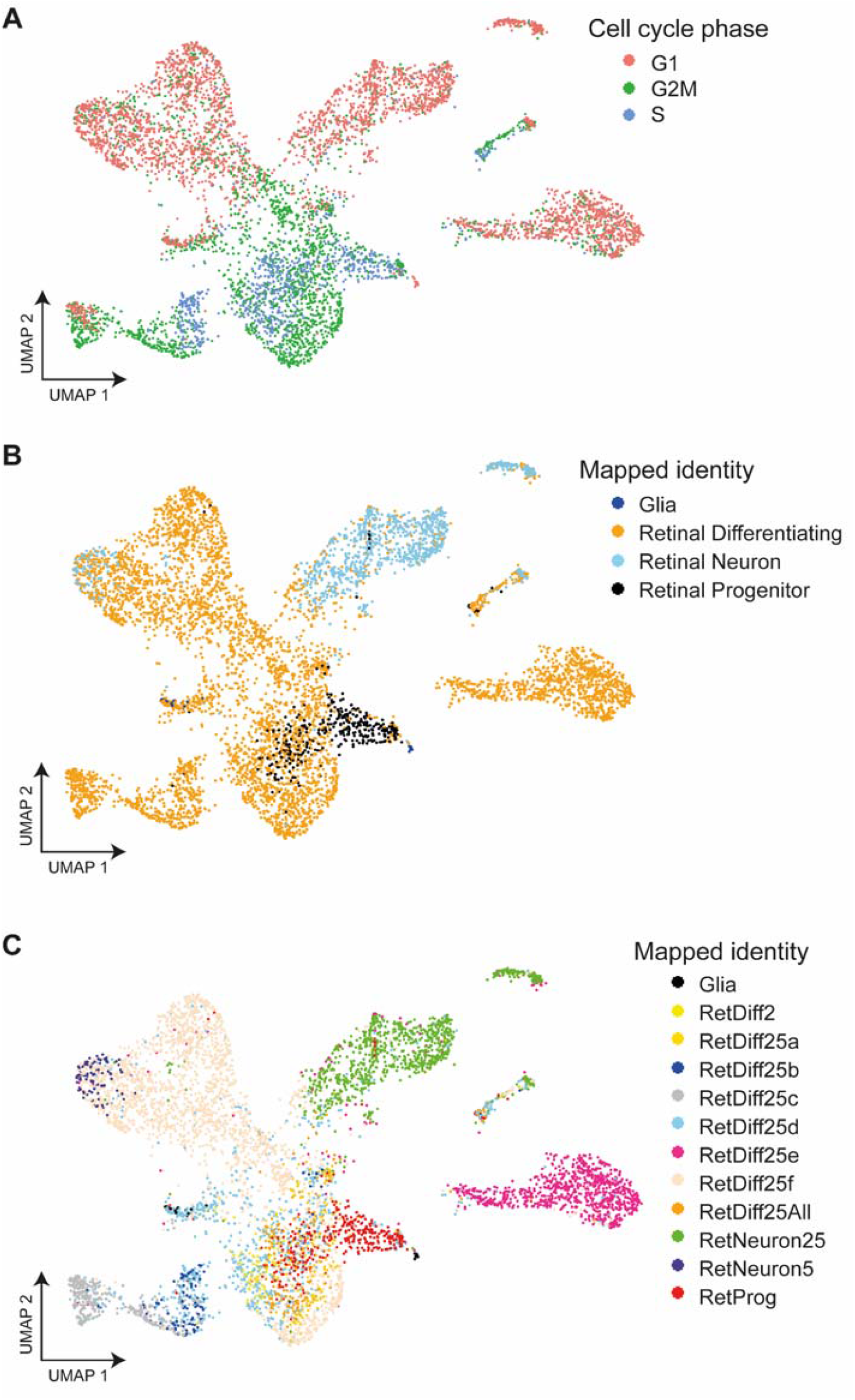
Mapped cell type identities and assigned cell cycle phase. **(A)** Cell cycle phase assigned to cells using Seurat visualised on a UMAP plot. **(B)** Simplified identity of cells mapped to the Farnsworth et al. (2019) single-cell expression zebrafish atlas (2 dpf), visualised on a UMAP plot; **(C)** shows actual mapped sub-types for retinal differentiating and neuron identities.

**Supplementary figure 4.**
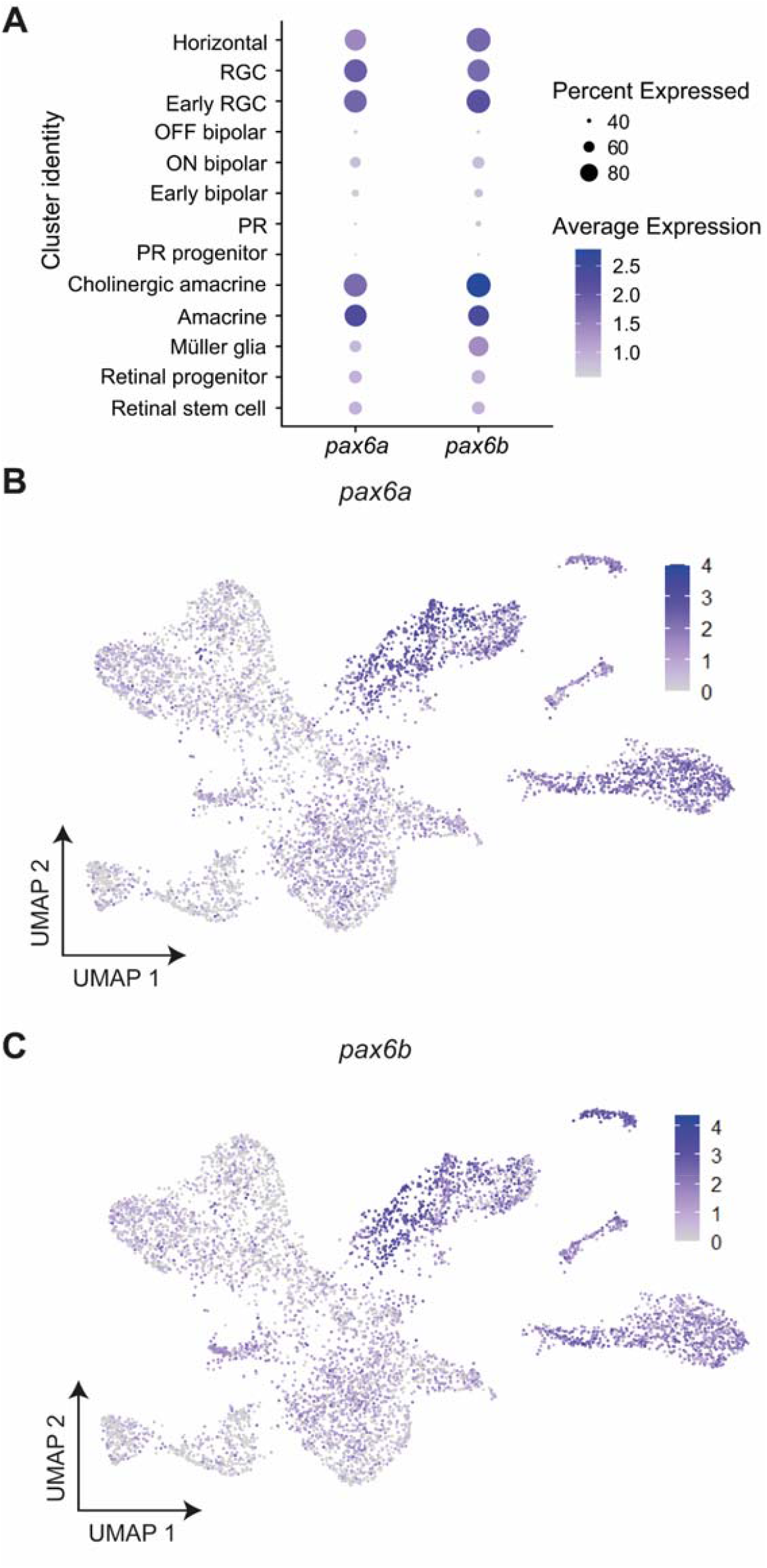
Expression of *pax6a* and *pax6b* across retinal cell types. **(A)** Dot plot showing average expression and percentage of cells expressing *pax6a* and *pax6b* in cell type clusters. **(B)** Expression of *pax6a* and **(C)** *pax6b* in cells visualised on UMAP plots.

**Supplementary figure 5.**
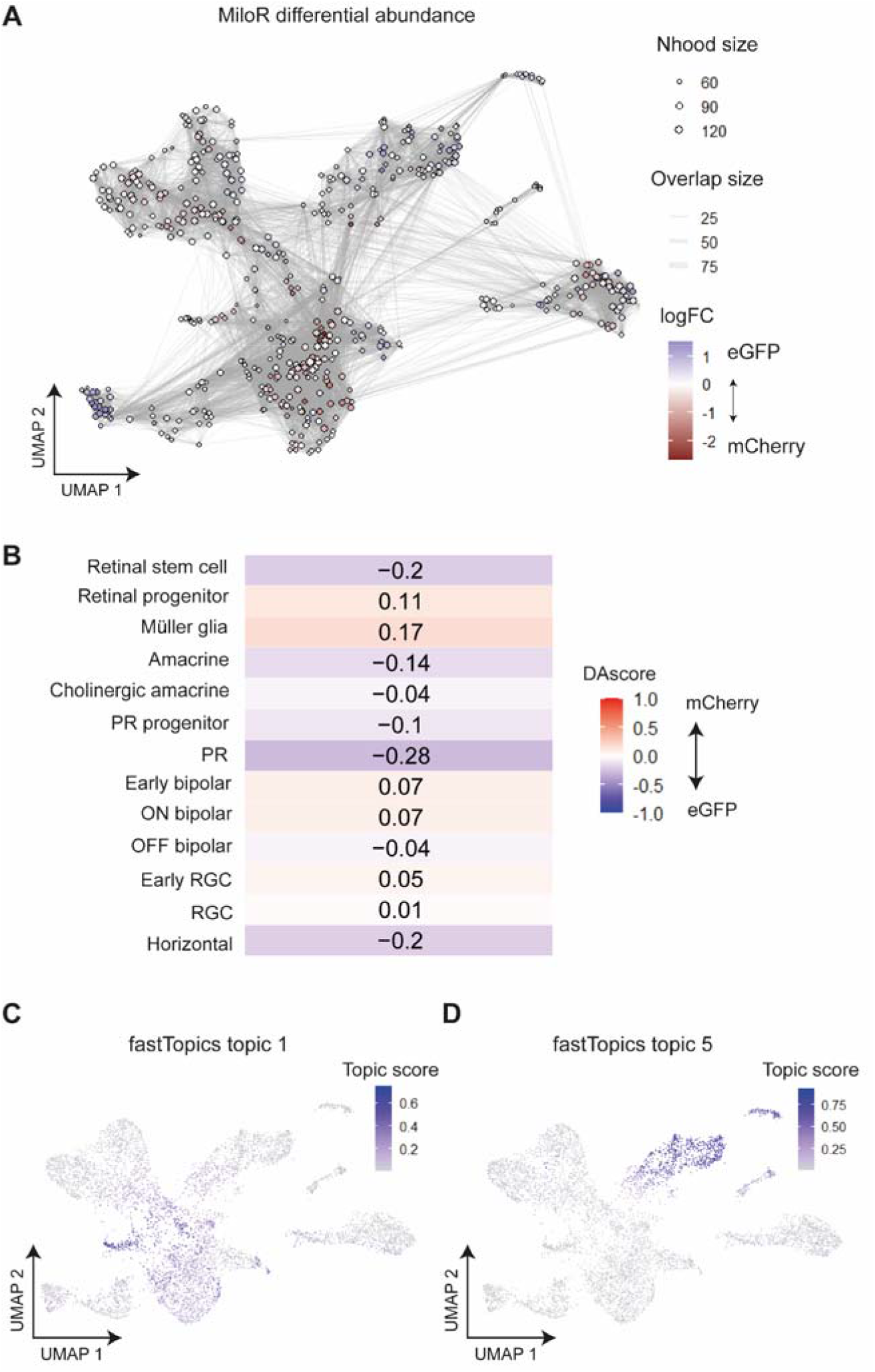
Differential abundance analysis and topic modelling validates the assignment of enhancer-active cell types. **(A)** MiloR (Dann et al., 2022) differential abundance testing. MiloR groups cells into neighbourhoods (nodes), then tests for differential abundance of cells from mCherry (HS5)-enriched or eGFP (NRE)-enriched samples within each neighbourhood. Neighbourhoods are shown here on a UMAP plot coloured according to the logFC score representing the abundance of cells from eGFP-enriched samples (positive, blue) or mCherry-enriched samples (negative, red) within each neighbourhood. **(B)** DAseq scores per cluster. Red (positive scores) indicates enrichment of cells from mCherry samples, blue (negative scores) for cells from eGFP samples. **(C)** fastTopics (Dey et al., 2017) applies topic modelling to count data. Each topic is defined by a set of genes and each cell is represented as a unique mixture of all topics. The score or ‘membership’ of each cell to topic 1 is shown here on a UMAP plot. Topic 2 appears to score highly in cells from Müller glia and progenitor clusters. **(D)** As in **(C)** showing the scores for topic 5, which are high in amacrine cell clusters.

**Supplementary movie 1. Time-lapse imaging reveals NRE and HS5 activity during development**.

Time-lapse imaging of an NRE-eGFP/HS5-mCherry F1 embryo from 24-70 hpf. Scale bar 100 μm.

## References

Angueyra JM, Kindt KS. 2018. Leveraging Zebrafish to Study Retinal Degenerations. Frontiers in Cell and Developmental Biology 6.

Aran D, Looney AP, Liu L, Wu E, Fong V, Hsu A, Chak S, Naikawadi RP, Wolters PJ, Abate AR, Butte AJ, Bhattacharya M. 2019. Reference-based analysis of lung single-cell sequencing reveals a transitional profibrotic macrophage. Nat Immunol 20:163–172. doi:10.1038/s41590-018-0276-y

Bhatia S, Bengani H, Fish M, Brown A, Divizia MT, De Marco R, Damante G, Grainger R, Van Heyningen V, Kleinjan DA. 2013. Disruption of autoregulatory feedback by a mutation in a remote, ultraconserved PAX6 enhancer causes aniridia. American Journal of Human Genetics 93:1126–1134. doi:10.1016/j.ajhg.2013.10.028

Bhatia S, Kleinjan DJ, Uttley K, Mann A, Dellepiane N, Bickmore WA. 2021. Quantitative spatial and temporal assessment of regulatory element activity in zebrafish. eLife 10. doi:10.7554/ELIFE.65601

Bhatia S, Monahan J, Ravi V, Gautier P, Murdoch E, Brenner S, Van Heyningen V, Venkatesh B, Kleinjan DA. 2014. A survey of ancient conserved non-coding elements in the PAX6 locus reveals a landscape of interdigitated cis-regulatory archipelagos. Developmental Biology 387:214–228. doi:10.1016/j.ydbio.2014.01.007

Butler A, Hoffman P, Smibert P, Papalexi E, Satija R. 2018. Integrating single-cell transcriptomic data across different conditions, technologies, and species. Nat Biotechnol 36:411–420. doi:10.1038/nbt.4096

Consortium TEP. 2012. An integrated encyclopedia of DNA elements in the human genome. Nature 489:57–74. doi:10.1038/nature11247

Dann E, Henderson NC, Teichmann SA, Morgan MD, Marioni JC. 2022. Differential abundance testing on single-cell data using k-nearest neighbor graphs. Nat Biotechnol 40:245–253. doi:10.1038/s41587-021-01033-z

Dey KK, Hsiao CJ, Stephens M. 2017. Visualizing the structure of RNA-seq expression data using grade of membership models. PLOS Genetics 13:e1006599. doi:10.1371/journal.pgen.1006599

Dickel DE, Ypsilanti AR, Pla R, Zhu Y, Barozzi I, Mannion BJ, Khin YS, Fukuda-Yuzawa Y, Plajzer-Frick I, Pickle CS, Lee EA, Harrington AN, Pham QT, Garvin TH, Kato M, Osterwalder M, Akiyama JA, Afzal V, Rubenstein JLR, Pennacchio LA, Visel A. 2018. Ultraconserved Enhancers Are Required for Normal Development. Cell 172:491-499.e15. doi:10.1016/j.cell.2017.12.017

Dupacova N, Antosova B, Paces J, Kozmik Z. 2021. Meis homeobox genes control progenitor competence in the retina. Proceedings of the National Academy of Sciences of the United States of America 118. doi:10.1073/PNAS.2013136118/SUPPL_FILE/PNAS.2013136118.SD01.XLSX

Ekström P, Johansson K. 2003. Differentiation of ganglion cells and amacrine cells in the rat retina: correlation with expression of HuC/D and GAP-43 proteins. Developmental Brain Research 145:1–8. doi:10.1016/S0165-3806(03)00170-6

Farhy C, Elgart M, Shapira Z, Oron-Karni V, Yaron O, Menuchin Y, Rechavi G, Ashery-Padan R. 2013. Pax6 Is Required for Normal Cell-Cycle Exit and the Differentiation Kinetics of Retinal Progenitor Cells. PLoS One 8:e76489. doi:10.1371/journal.pone.0076489

Farley EK, Olson KM, Zhang W, Brandt AJ, Rokhsar DS, Levine MS. 2015. Suboptimization of developmental enhancers. Science 350:325–8. doi:10.1126/science.aac6948

Farley EK, Olson KM, Zhang W, Rokhsar DS, Levine MS. 2016. Syntax compensates for poor binding sites to encode tissue specificity of developmental enhancers. Proceedings of the National Academy of Sciences 113:6508–6513. doi:10.1073/pnas.1605085113

Farnsworth DR, Saunders LM, Miller AC. 2019. A single-cell transcriptome atlas for zebrafish development. Developmental Biology. doi:10.1016/j.ydbio.2019.11.008

Fischer AJ, Bosse JL, El-Hodiri HM. 2013. The ciliary marginal zone (CMZ) in development and regeneration of the vertebrate eye. Experimental Eye Research 116:199–204. doi:10.1016/j.exer.2013.08.018

Goldman D. 2014. Müller glia cell reprogramming and retina regeneration. Nat Rev Neurosci 15:431–442. doi:10.1038/nrn3723

Griffin C, Kleinjan DA, Doe B, van Heyningen V. 2002. New 31. elements control Pax6 expression in the developing pretectum, neural retina and olfactory region. Mechanisms of Development 112:89–100. doi:10.1016/S0925-4773(01)00646-3

Halder G, Callaerts P, Gehring WJ. 1995. Induction of Ectopic Eyes by Targeted Expression of the eyeless Gene in Drosophila. Science 267:1788–1792. doi:10.1126/science.7892602

Jadhav AP, Roesch K, Cepko CL. 2009. Development and neurogenic potential of Müller glial cells in the vertebrate retina. Progress in Retinal and eye Research 28:249–262. doi:10.1016/J.PRETEYERES.2009.05.002

Jindal GA, Farley EK. 2021. Enhancer grammar in development, evolution, and disease: dependencies and interplay. Developmental Cell 56:575–587. doi:10.1016/j.devcel.2021.02.016

Joly S, Pernet V, Samardzija M, Grimm C. 2011. Pax6-positive müller glia cells express cell cycle markers but do not proliferate after photoreceptor injury in the mouse retina. Glia 59:1033–1046. doi:10.1002/glia.21174

Kammandel B, Chowdhury K, Stoykova A, Aparicio S, Brenner S, Gruss P. 1999. Distinct cis-essential modules direct the time-space pattern of the Pax6 gene activity. Developmental Biology 205:79–97. doi:10.1006/dbio.1998.9128

Kharchenko PV, Silberstein L, Scadden DT. 2014. Bayesian approach to single-cell differential expression analysis. Nat Methods 11:740–742. doi:10.1038/nmeth.2967

Kim Y, Lim S, Ha T, Song YH, Sohn YI, Park DJ, Paik SS, Kim-Kaneyama JR, Song MR, Leung A, Levine EM, Kim IB, Goo YS, Lee SH, Kang KH, Kim JW. 2017. The LIM protein complex establishes a retinal circuitry of visual adaptation by regulating Pax6 α-enhancer activity. eLife 6. doi:10.7554/ELIFE.21303

Kimmel CB, Ballard WW, Kimmel SR, Ullmann B, Schilling TF. 1995. Stages of embryonic development of the zebrafish. Developmental Dynamics 203:253–310. doi:10.1002/aja.1002030302

Kleinjan DA, Seawright A, Schedl A, Quinlan RA, Danes S, van Heyningen V. 2001. Aniridia-associated translocations, DNase hypersensitivity, sequence comparison and transgenic analysis redefine the functional domain of PAX6. Human Molecular Genetics 10:2049–2059. doi:10.1093/hmg/10.19.2049

Korsunsky I, Millard N, Fan J, Slowikowski K, Zhang F, Wei K, Baglaenko Y, Brenner M, Loh P, Raychaudhuri S. 2019. Fast, sensitive and accurate integration of single-cell data with Harmony. Nat Methods 16:1289–1296. doi:10.1038/s41592-019-0619-0

Kvon EZ, Waymack R, Elabd MG, Wunderlich Z. 2021. Enhancer redundancy in development and disease. Nature Reviews Genetics 1–13. doi:10.1038/s41576-020-00311-x

Kvon EZ, Zhu Y, Kelman G, Novak CS, Plajzer-Frick I, Kato M, Garvin TH, Pham Q, Harrington AN, Hunter RD, Godoy J, Meky EM, Akiyama JA, Afzal V, Tran S, Escande F, Gilbert-Dussardier B, Jean-Marçais N, Hudaiberdiev S, Ovcharenko I, Dobbs MB, Gurnett CA, Manouvrier-Hanu S, Petit F, Visel A, Dickel DE, Pennacchio LA. 2020. Comprehensive In Vivo Interrogation Reveals Phenotypic Impact of Human Enhancer Variants. Cell 180:1262-1271.e15. doi:10.1016/j.cell.2020.02.031

Lauderdale JD, Wilensky JS, Oliver ER, Walton DS, Glaser T. 2000. 3’ deletions cause aniridia by preventing PAX6 gene expression. Proceedings of the National Academy of Sciences 97:13755–13759. doi:10.1073/pnas.240398797

Lettice LA, Heaney SJH, Purdie LA, Li L, de Beer P, Oostra BA, Goode D, Elgar G, Hill RE, de Graaff E. 2003. A long-range Shh enhancer regulates expression in the developing limb and fin and is associated with preaxial polydactyly. Human Molecular Genetics 12:1725–1735. doi:10.1093/hmg/ddg180

Li S, Mo Z, Yang X, Price SM, Shen MM, Xiang M. 2004. Foxn4 Controls the Genesis of Amacrine and Horizontal Cells by Retinal Progenitors. Neuron 43:795–807. doi:10.1016/j.neuron.2004.08.041

Lim F, Ryan GE, L. SH, Solvason JJ, Steffen P, Farley EK. 2022. Affinity-optimizing variants within the ZRS enhancer disrupt limb development. bioRxiv 2022.05.27.493789. doi:10.1101/2022.05.27.493789

Lima Cunha D, Arno G, Corton M, Moosajee M. 2019. The Spectrum of PAX6 Mutations and Genotype-Phenotype Correlations in the Eye. Genes 10:1050. doi:10.3390/genes10121050

Long HK, Osterwalder M, Welsh IC, Hansen K, Davies JOJ, Liu YE, Koska M, Adams AT, Aho R, Arora N, Ikeda K, Williams RM, Sauka-Spengler T, Porteus MH, Mohun T, Dickel DE, Swigut T, Hughes JR, Higgs DR, Visel A, Selleri L, Wysocka J. 2020. Loss of Extreme Long-Range Enhancers in Human Neural Crest Drives a Craniofacial Disorder. Cell Stem Cell 27. doi:10.1016/j.stem.2020.09.001

Long HK, Prescott SL, Wysocka J. 2016. Ever-Changing Landscapes: Transcriptional Enhancers in Development and Evolution. Cell 167:1170–1187. doi:10.1016/J.CELL.2016.09.018

Lun ATL, Riesenfeld S, Andrews T, Dao TP, Gomes T, Marioni JC, participants in the 1st Human Cell Atlas Jamboree. 2019. EmptyDrops: distinguishing cells from empty droplets in droplet-based single-cell RNA sequencing data. Genome Biology 20:63. doi:10.1186/s13059-019-1662-y

Marquardt T, Ashery-Padan R, Andrejewski N, Scardigli R, Guillemot F, Gruss P. 2001. Pax6 Is Required for the Multipotent State of Retinal Progenitor Cells. Cell 105:43–55. doi:10.1016/S0092-8674(01)00295-1

Maurano MT, Humbert R, Rynes E, Thurman RE, Haugen E, Wang H, Reynolds AP, Sandstrom R, Qu H, Brody J, Shafer A, Neri F, Lee K, Kutyavin T, Stehling-Sun S, Johnson AK, Canfield TK, Giste E, Diegel M, Bates D, Hansen RS, Neph S, Sabo PJ, Heimfeld S, Raubitschek A, Ziegler S, Cotsapas C, Sotoodehnia N, Glass I, Sunyaev SR, Kaul R, Stamatoyannopoulos JA. 2012. Systematic localization of common disease-associated variation in regulatory DNA. Science 337:1190–5. doi:10.1126/science.1222794

McBride DJ, Buckle A, van Heyningen V, Kleinjan DA. 2011. DNaseI Hypersensitivity and Ultraconservation Reveal Novel, Interdependent Long-Range Enhancers at the Complex Pax6 Cis-Regulatory Region. PLoS ONE 6:e28616. doi:10.1371/journal.pone.0028616

McCarthy DJ, Campbell KR, Lun ATL, Wills QF. 2017. Scater: pre-processing, quality control, normalization and visualization of single-cell RNA-seq data in R. Bioinformatics 33:1179–1186. doi:10.1093/bioinformatics/btw777

Meuleman W, Muratov A, Rynes E, Halow J, Lee K, Bates D, Diegel M, Dunn D, Neri F, Teodosiadis A, Reynolds A, Haugen E, Nelson J, Johnson A, Frerker M, Buckley M, Sandstrom R, Vierstra J, Kaul R, Stamatoyannopoulos J. 2020. Index and biological spectrum of human DNase I hypersensitive sites. Nature 584:244–251. doi:10.1038/s41586-020-2559-3

Onuma Y, Takahashi S, Asashima M, Kurata S, Gehring WJ. 2002. Conservation of Pax 6 function and upstream activation by Notch signaling in eye development of frogs and flies. Proceedings of the National Academy of Sciences 99:2020–2025. doi:10.1073/pnas.022626999

Osterwalder M, Barozzi I, Tissières V, Fukuda-Yuzawa Y, Mannion BJ, Afzal SY, Lee EA, Zhu Y, Plajzer-Frick I, Pickle CS, Kato M, Garvin TH, Pham QT, Harrington AN, Akiyama JA, Afzal V, Lopez-Rios J, Dickel DE, Visel A, Pennacchio LA. 2018. Enhancer redundancy provides phenotypic robustness in mammalian development. Nature 554:239–243. doi:10.1038/nature25461

Plaza S, Dozier C, Langlois MC, Saule S. 1995. Identification and characterization of a neuroretina-specific enhancer element in the quail Pax-6 (Pax-QNR) gene. Molecular and Cellular Biology 15:892–903. doi:10.1128/mcb.15.2.892

R Core Team. 2021. R: A Language and Environment for Statistical Computing.

Ravi V, Bhatia S, Gautier P, Loosli F, Tay BH, Tay A, Murdoch E, Coutinho P, van Heyningen V, Brenner S, Venkatesh B, Kleinjan DA. 2013. Sequencing of Pax6 Loci from the Elephant Shark Reveals a Family of Pax6 Genes in Vertebrate Genomes, Forged by Ancient Duplications and Divergences. PLoS Genetics 9:e1003177. doi:10.1371/journal.pgen.1003177

Ravi V, Bhatia S, Shingate P, Tay BH, Venkatesh B, Kleinjan DA. 2019. Lampreys, the jawless vertebrates, contain three Pax6 genes with distinct expression in eye, brain and pancreas. Scientific Reports 9:1–12. doi:10.1038/s41598-019-56085-8

Remez LA, Onishi A, Menuchin-Lasowski Y, Biran A, Blackshaw S, Wahlin KJ, Zack DJ, Ashery-Padan R. 2017. Pax6 is essential for the generation of late-born retinal neurons and for inhibition of photoreceptor-fate during late stages of retinogenesis. Developmental Biology 432:140–150. doi:10.1016/j.ydbio.2017.09.030

Richardson R, Tracey-White D, Webster A, Moosajee M. 2016. The zebrafish eye—a paradigm for investigating human ocular genetics. Eye 2017 31:1 31:68–86. doi:10.1038/eye.2016.198

Rulands S, Iglesias-Gonzalez AB, Boije H. 2018. Deterministic fate assignment of Müller glia cells in the zebrafish retina suggests a clonal backbone during development. The European Journal of Neuroscience 48:3597. doi:10.1111/EJN.14257

Schindelin J, Arganda-Carreras I, Frise E, Kaynig V, Longair M, Pietzsch T, Preibisch S, Rueden C, Saalfeld S, Schmid B, Tinevez J-Y, White DJ, Hartenstein V, Eliceiri K, Tomancak P, Cardona A. 2012. Fiji: an open-source platform for biological-image analysis. Nat Methods 9:676–682. doi:10.1038/nmeth.2019

Schmitt EA, Dowling JE. 1999. Early retinal development in the zebrafish, Danio rerio: light and electron microscopic analyses. J Comp Neurol 404:515–536.

Shi X, Shin Teo L, Pan X, Chong S-W, Kraut R, Korzh V, Wohland T. 2009. Probing events with single molecule sensitivity in zebrafish and Drosophila embryos by fluorescence correlation spectroscopy. Developmental Dynamics 238:3156–3167. doi:10.1002/dvdy.22140

Singhal S, Bhatia B, Jayaram H, Becker S, Jones MF, Cottrill PB, Khaw PT, Salt TE, Limb GA. 2012. Human Müller Glia with Stem Cell Characteristics Differentiate into Retinal Ganglion Cell (RGC) Precursors In Vitro and Partially Restore RGC Function In Vivo Following Transplantation. Stem Cells Transl Med 1:188–199. doi:10.5966/sctm.2011-0005

Smemo S, Campos LC, Moskowitz IP, Krieger JE, Pereira AC, Nobrega MA. 2012. Regulatory variation in a TBX5 enhancer leads to isolated congenital heart disease. Human molecular genetics 21:3255–63. doi:10.1093/hmg/dds165

Snetkova V, Ypsilanti AR, Akiyama JA, Mannion BJ, Plajzer-Frick I, Novak CS, Harrington AN, Pham QT, Kato M, Zhu Y, Godoy J, Meky E, Hunter RD, Shi M, Kvon EZ, Afzal V, Tran S, Rubenstein JLR, Visel A, Pennacchio LA, Dickel DE. 2021. Ultraconserved enhancer function does not require perfect sequence conservation. Nature Genetics 1–8. doi:10.1038/s41588-021-00812-3

Sprague J, Bayraktaroglu L, Bradford Y, Conlin T, Dunn N, Fashena D, Frazer K, Haendel M, Howe DG, Knight J, Mani P, Moxon SAT, Pich C, Ramachandran S, Schaper K, Segerdell E, Shao X, Singer A, Song P, Sprunger B, Van Slyke CE, Westerfield M. 2008. The Zebrafish Information Network: the zebrafish model organism database provides expanded support for genotypes and phenotypes. Nucleic Acids Res 36:D768–D772. doi:10.1093/nar/gkm956

Suzuki-Kerr H, Iwagawa T, Sagara H, Mizota A, Suzuki Y, Watanabe S. 2018. Pivotal roles of Fezf2 in differentiation of cone OFF bipolar cells and functional maturation of cone ON bipolar cells in retina. Experimental Eye Research 171:142–154. doi:10.1016/j.exer.2018.03.017

Thummel R, Kassen SC, Enright JM, Nelson CM, Montgomery JE, Hyde DR. 2008. Characterization of Müller glia and neuronal progenitors during adult zebrafish retinal regeneration. Exp Eye Res 87:433–444. doi:10.1016/j.exer.2008.07.009

Tyas DA, Simpson TI, Carr CB, Kleinjan DA, van Heyningen V, Mason JO, Price DJ. 2006. Functional conservation of Pax6 regulatory elements in humans and mice demonstrated with a novel transgenic reporter mouse. BMC Dev Biol 6:21. doi:10.1186/1471-213X-6-21

van Beest EH, Mukherjee S, Kirchberger L, Schnabel UH, van der Togt C, Teeuwen RRM, Barsegyan A, Meyer AF, Poort J, Roelfsema PR, Self MW. 2021. Mouse visual cortex contains a region of enhanced spatial resolution. Nature Communications 2021 12:1 12:1–17. doi:10.1038/s41467-021-24311-5

van Heyningen V. 2002. PAX6 in sensory development. Human Molecular Genetics 11:1161–1167. doi:10.1093/hmg/11.10.1161

Wan Y, Almeida AD, Rulands S, Chalour N, Muresan L, Wu Y, Simons BD, He J, Harris WA. 2016. The ciliary marginal zone of the zebrafish retina: clonal and time-lapse analysis of a continuously growing tissue. Development 143:1099–1107. doi:10.1242/dev.133314

Watanabe-Susaki K, Takada H, Enomoto K, Miwata K, Ishimine H, Intoh A, Ohtaka M, Nakanishi M, Sugino H, Asashima M, Kurisaki A. 2014. Biosynthesis of Ribosomal RNA in Nucleoli Regulates Pluripotency and Differentiation Ability of Pluripotent Stem Cells. Stem Cells 32:3099–3111. doi:10.1002/stem.1825

Whitney IE, Keeley PW, St. John AJ, Kautzman AG, Kay JN, Reese BE. 2014. Sox2 Regulates Cholinergic Amacrine Cell Positioning and Dendritic Stratification in the Retina. J Neurosci 34:10109–10121. doi:10.1523/JNEUROSCI.0415-14.2014

Wickham H. 2009. ggplot2: Elegant Graphics for Data Analysis, Use R! Springer New York, NY.

Williams SC, Altmann CR, Chow RL, Hemmati-Brivanlou A, Lang RA. 1998. A highly conserved lens transcriptional control element from the Pax-6 gene. Mechanisms of Development 73:225–229. doi:10.1016/S0925-4773(98)00057-4

Yoshimatsu T, Schröder C, Nevala NE, Berens P, Baden T. 2020. Fovea-like Photoreceptor Specializations Underlie Single UV Cone Driven Prey-Capture Behavior in Zebrafish. Neuron 107:320. doi:10.1016/J.NEURON.2020.04.021

Zhao J, Jaffe A, Li H, Lindenbaum O, Sefik E, Jackson R, Cheng X, Flavell RA, Kluger Y. 2021. Detection of differentially abundant cell subpopulations in scRNA-seq data. Proceedings of the National Academy of Sciences 118:e2100293118. doi:10.1073/pnas.2100293118

Zheng GXY, Terry JM, Belgrader P, Ryvkin P, Bent ZW, Wilson R, Ziraldo SB, Wheeler TD, McDermott GP, Zhu J, Gregory MT, Shuga J, Montesclaros L, Underwood JG, Masquelier DA, Nishimura SY, Schnall-Levin M, Wyatt PW, Hindson CM, Bharadwaj R, Wong A, Ness KD, Beppu LW, Deeg HJ, McFarland C, Loeb KR, Valente WJ, Ericson NG, Stevens EA, Radich JP, Mikkelsen TS, Hindson BJ, Bielas JH. 2017. Massively parallel digital transcriptional profiling of single cells. Nat Commun 8:14049. doi:10.1038/ncomms14049

