## Supplementary figures and images for "Unique functions of two overlapping *PAX6* retinal enhancers"

### Supplementary figure 1

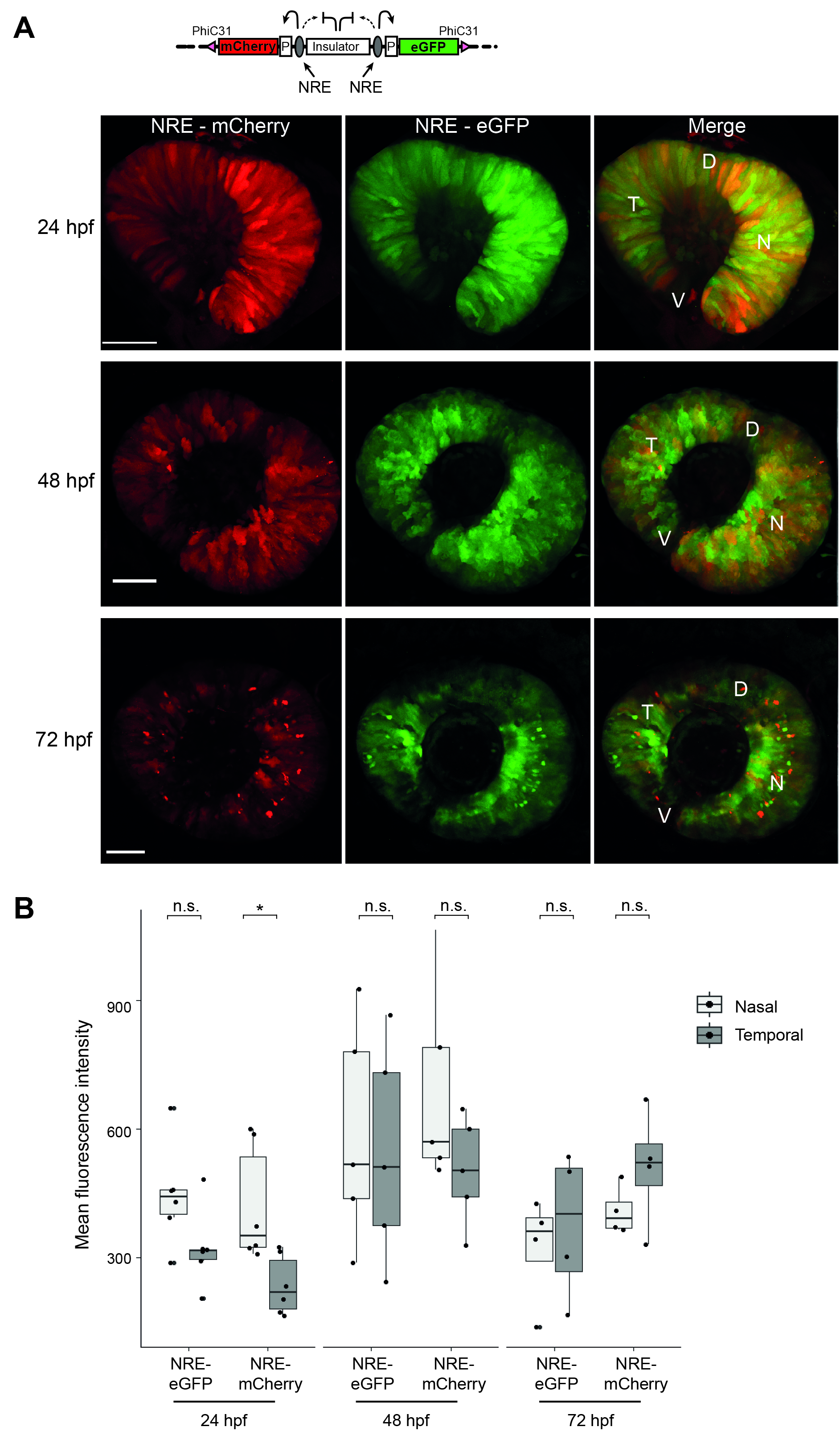

### Supplementary figure 2

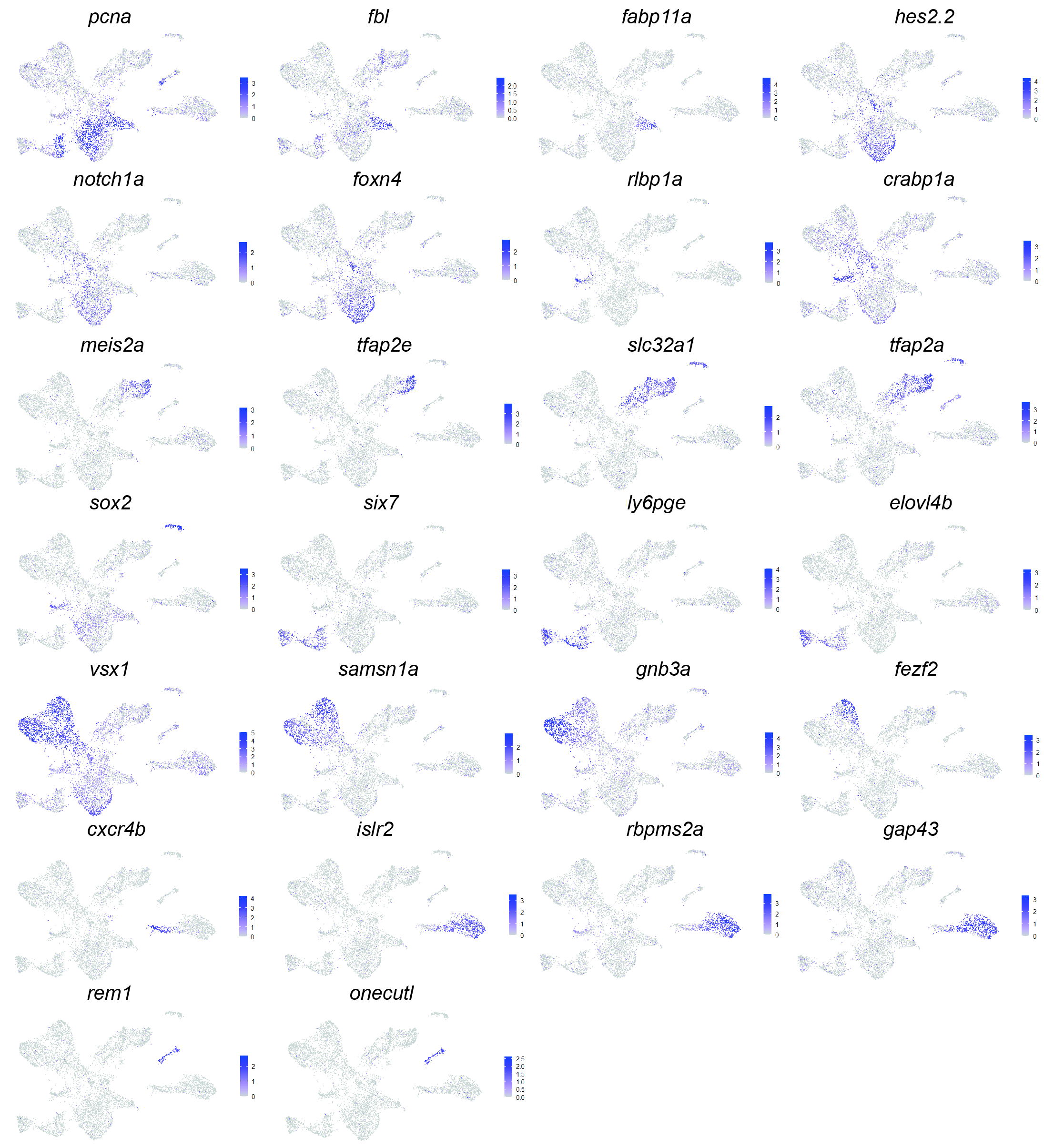

### Supplementary figure 3

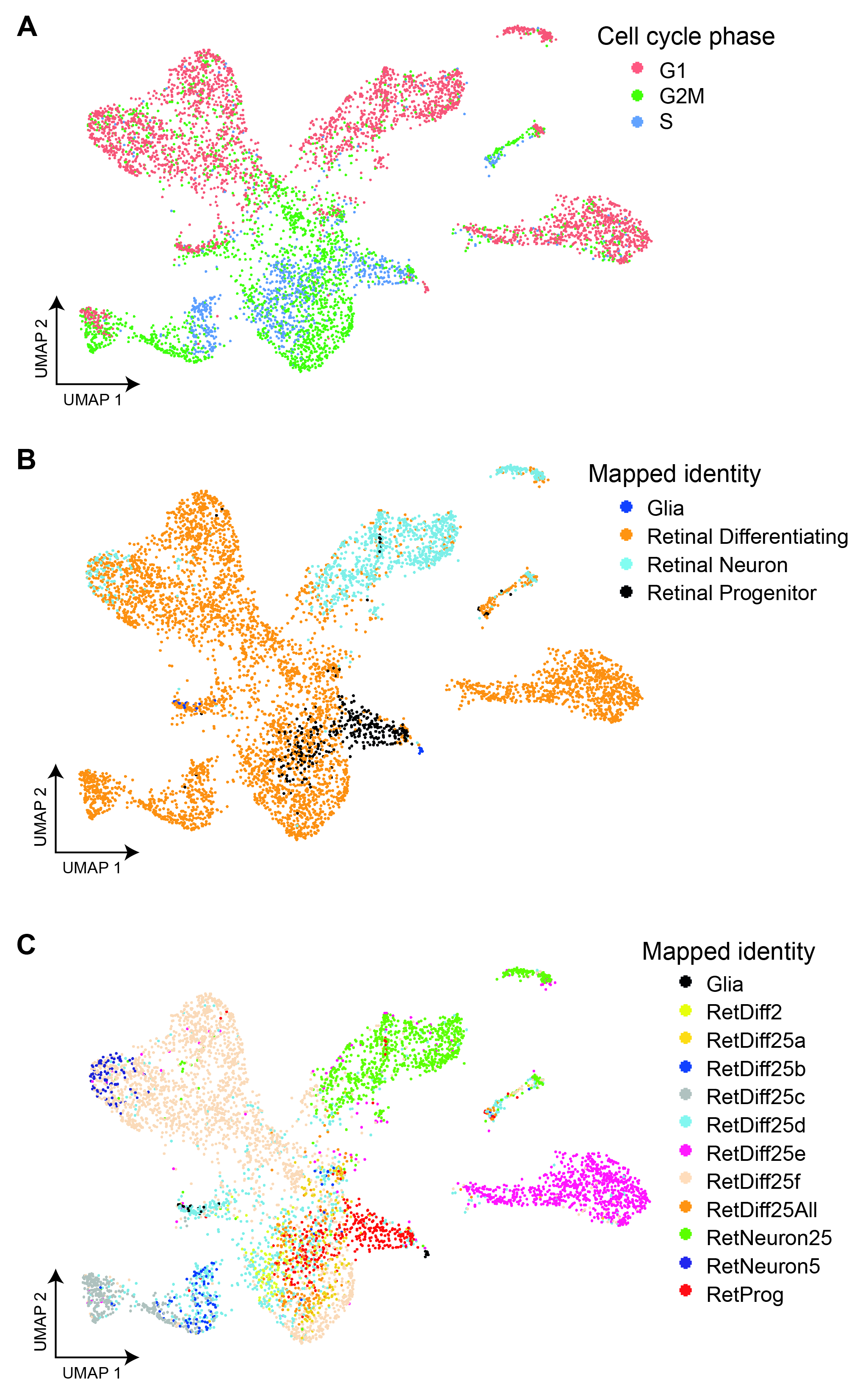

### Supplementary figure 4

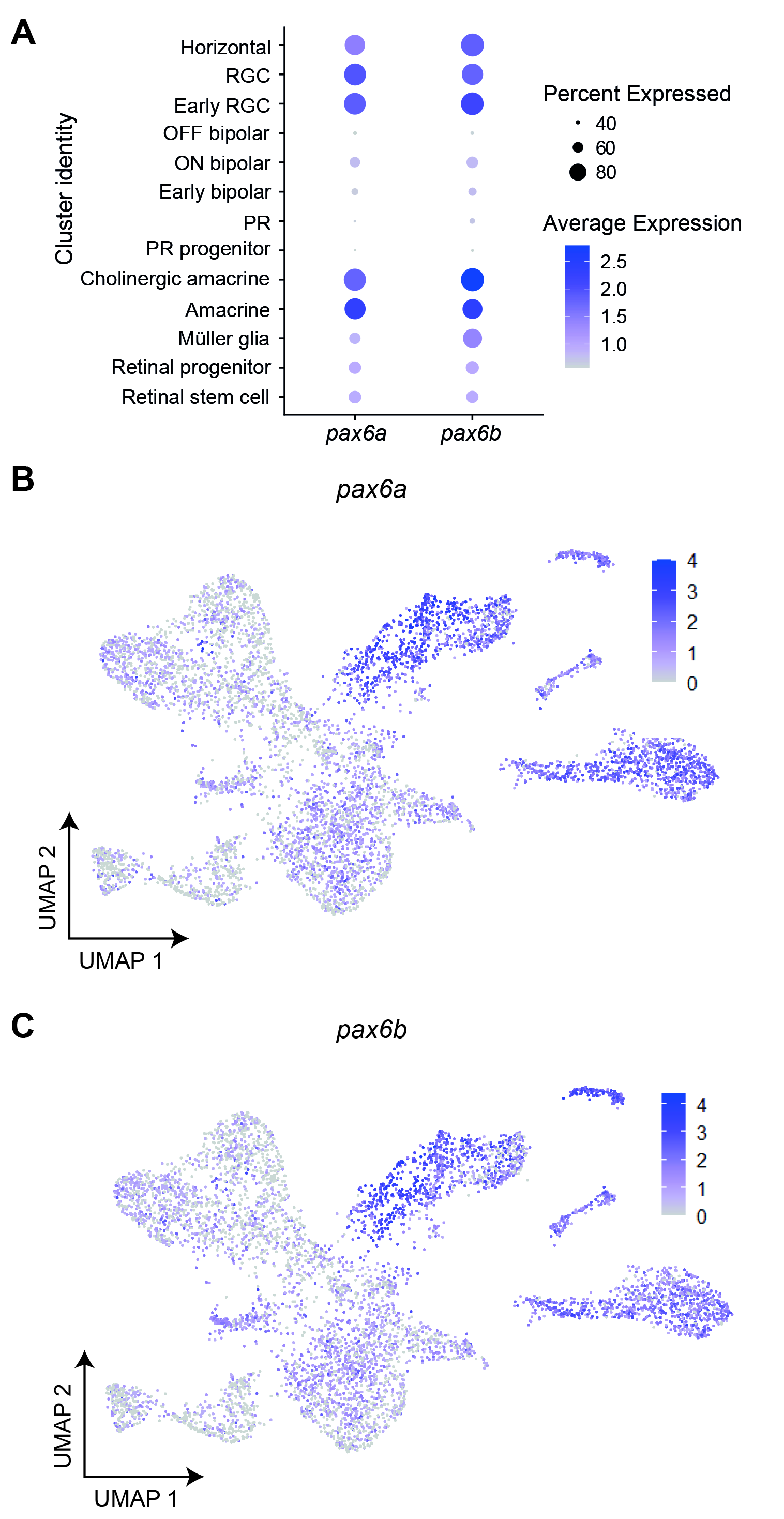

### Supplementary figure 5

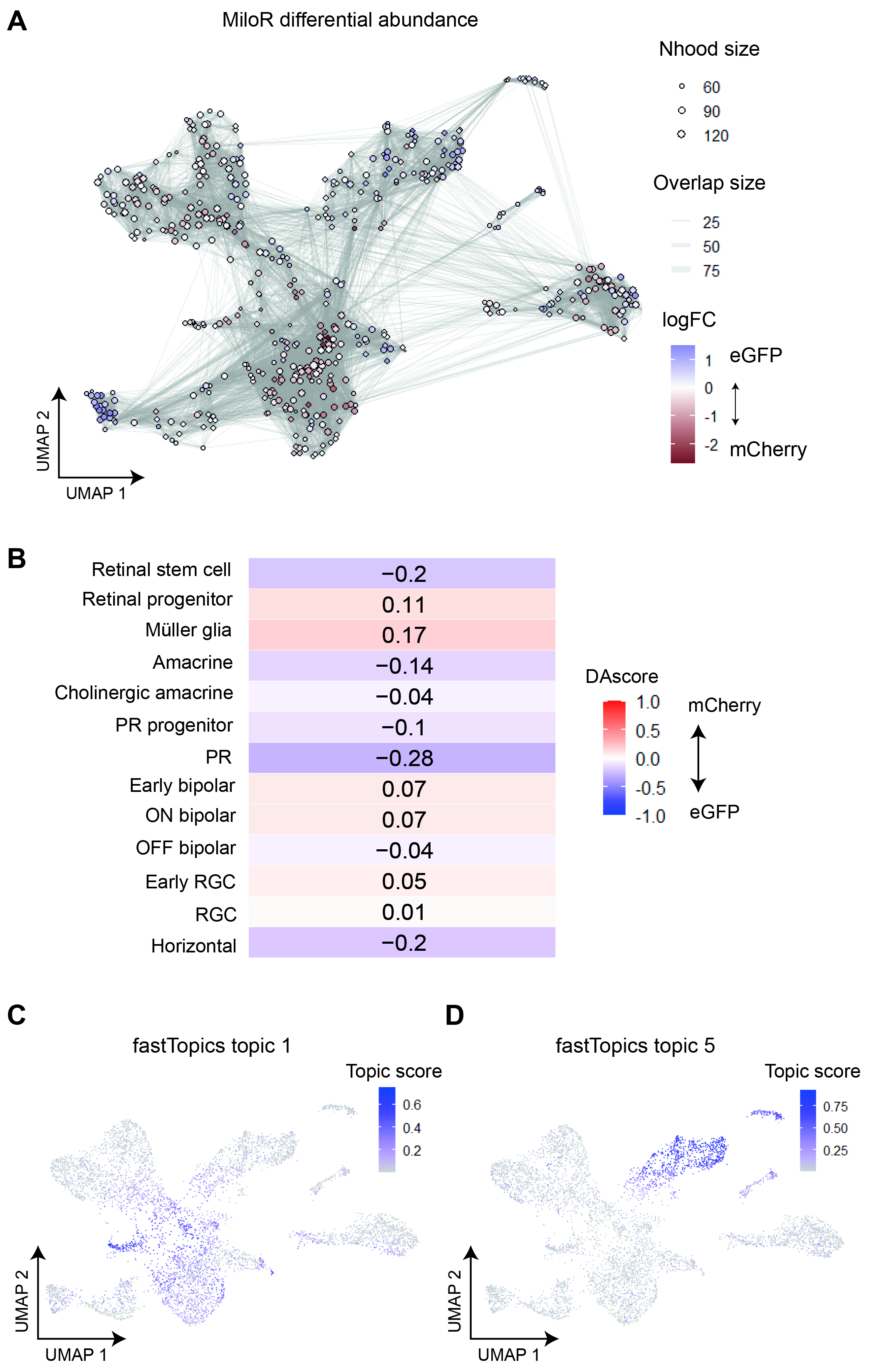
